# Introgression across ploidies contributes to genetic diversity in introduced urban *Capsella bursa-pastoris*

**DOI:** 10.64898/2026.03.17.712489

**Authors:** Maya K. Wilson Brown, Rebecca Panko, Adrian E. Platts, Emily B. Josephs

## Abstract

Successful establishment of a species in a new range is a useful way to understand the impact of demography and selection on the evolution of globally distributed species. In particular, introductions influence genetic diversity and population structure in the introduced range in unpredictable ways. Additionally, introgressive hybridization is often associated with successful establishment in new ranges. In this study, we explore the impact of introgressive hybridization on the polyploid *Capsella bursa-pastoris* in the New York City metropolitan area. We find *Capsella bursa-pastoris* in the New York City metropolitan area likely originated from multiple introductions from northern Eurasia, and that populations across the New York City metropolitan area are generally panmictic. As with *Capsella bursa-pastoris* in Eurasia, we discover evidence of introgression from the diploid *Capsella rubella* in this population. By evaluating ancestry in regions across the genome, we find introgressed regions are rich in gene content and contribute to genetic diversity in this population. These results suggest that introgressive hybridization before introductions may buffer species from the negative effects of population bottlenecks and allow for successful establishment.

**Significance:** Gene flow between groups of different ploidies is known to occur in natural systems, but has been understudied in evolutionary genetics until recently. In this study, we show that an introduced population of the allotetraploid *Capsella bursa-pastoris* contains introgressed material from the diploid *C. rubella.* The introgressed regions of the genome have a high degree of diversity suggesting that introgression from *C. rubella* to *C. bursa-pastoris* may have contributed to the overall high degree of diversity in the species. Understanding the evolutionary impact of gene flow between ploidies is an important step to identifying what makes some polyploid lineages successful.

## Introduction

Studying range expansions can reveal how evolutionary forces impact species success. Populations that are introduced to new habitats beyond their natural dispersal range provide semi-natural experiments to explore the consequences of introduction on genetic population structure and diversity (Huey et al. 2005; Whitney and Gabler 2008; Colautti and Lau 2016). Understanding the genetic structure of introduced populations, along with identifying their source populations, provides crucial insight to the demographic history of a species introduction such as founding events and the number of introductions, as well as interactions post-introduction (Dlugosch and Parker 2008; Le Roux and Wieczorek 2009). The genetic structure of introduced populations also influences genetic diversity, since non-random mating reduces the effective population size, altering the effects of selection and drift. Moreover, we expect genetic diversity to decrease in introduced ranges due to population bottlenecks and founder effects, but that is not always the case (Squirrell et al. 2001; Kolbe et al. 2004).

Introgressive hybridization, or introgression, is frequently associated with successful establishment of introduced species because it can efficiently introduce adaptative alleles and alter life-history traits (Rieseberg et al. 1999; Ellstrand and Schierenbeck 2000; Baack and Rieseberg 2007; Abbott et al. 2013). More generally, the uptake of additional genetic material that differs among populations within a species increases intraspecific genetic variation. However, we do not yet know how this variation impacts a species when introduced to a new environment.

Throughout the last several decades, research on the evolutionary impacts of introgression have focused on hybridization between diploid species but gene flow is not limited to homoploid hybrids. Observations of introgression between ploidies is common (Ramsey and Schemske 1998; Petit et al. 1999; Jørgensen et al. 2011; Zohren et al. 2016; Bartolić et al. 2024), and this introgression can have major phenotypic effects (Chapman and Abbott 2010; Arnold et al. 2015; Arnold et al. 2016). As in gene flow between diploids, introgression between ploidies has the potential to introduce adaptive variants (Schmickle & Yant 2021) and increase genetic variation (Petit et al. 1999). Moreover, the dynamics of introgressed genetic material may be more complex in polyploids due to their high degree of variation in dominance effects, mutation rates, and recombination (Brown et al. 2024). For example, in polyploids that exhibit disomic inheritance (i.e. homeologous chromosomes do not recombine with one another), the effect of deleterious recessive variants could effectively be masked by alleles fixed in the other pair of homeologous chromosomes (Conover and Wendel 2022). Therefore, interploidal introgression should be considered when investigating the influence of past hybridization on species range expansions and intraspecific diversity.

In this study, we explore if past interploid introgression contributes to the genetic diversity of an introduced population of an urban weed. *Capsella bursa-pastoris* is a cosmopolitan Brassicaceae introduced to the United States only a few hundred years ago (Neuffer and Hurka 1999). In its ancestral Eurasian range, *C. bursa-pastoris* shows substantial evidence of introgression from a diploid congener, *Capsella rubella* (Slotte et al. 2008; Han et al. 2015; Kryvokhyzha et al. 2019). However, we do not know if introgressed genotypes exist in the introduced range or to what extent introgression impacts genetic variation in the species. We densely sampled *C. bursa-pastoris* in the New York City Metropolitan area to gain insight on the introduction history and evolutionary dynamics of this species. We then examine the relationship of *C. bursa-pastoris* in the New York City Metropolitan area with genotypes in the ancestral range to identify potential source populations and characterize how introduction to the non-native range may have influenced this species. Finally, we determine if introduced *C. bursa-pastoris* also carries genetic material from pre-introduction hybridization and investigate the role of interspecific introgression on the genetic diversity of this species. We use these data to inform if interploidal introgression may have contributed to the success of this species in the introduced range. Together, hybridization between ploidies is likely an underappreciated mechanism to explain the prevalence of polyploid lineages in successful range expansions.

## Results

### Reference genome assembly

For this study, we generated a reference genome from *C. bursa-pastoris* from Yonkers, New York, USA. We obtained approximately 65 gigbases (about 90x depth for our genome size) of Pacific Biosciences HiFi long-read data. Our assembly generated 16 major contigs and four smaller contigs. We numbered our contigs in accordance with another *C. bursa-pastoris* assembly (Fiscus 2022) for continuity across research groups. Synteny comparison to another assembly (Fiscus 2022) suggested that at least one of the minor contigs represented the short arm of chromosome 1 and another a peri-telomeric fragment of chromosome 4. We made two joins to make our NYC assembly close to syntenic with the other *C. bursa-pastoris* assemblies, leaving only two short unplaced contigs. The total assembly size was just over 351 Mb. The last 15 Mb of Chromosome 9 are gene-poor but highly repeat-rich and apparently specific to this New York City line (Supplemental Figure 1). Based on the transition from normal levels of transposable elements (TEs) and genes to an extended TE-enriched, gene-depleted sequence, it appears that there is a fusion point of a B-chromosome to the end of Chromosome 9 that occurs around the 28,230,891 basepair. Beyond this location, there are abrupt changes in TE and gene density trends, coincident with a set of serially reiterated telomeric repeats, possibly marking the end of the original chromosome 9 sequence. BUSCO analysis of our genome assembly indicates we captured 99.6 percent of highly conserved genes in the Brassicales HMM gene model. Of the BUSCO groups, 4103 of 4311 (95%) were found exactly twice, and 4223 (98%) were found in more than one copy (Supplemental Text 1). Consequently, 97% of multi-copy genes were found exactly twice. At the time of assembly, Hifisam did not have an option to explicitly preserve telomeric sequences. Nonetheless, 7 chromosomes had telomeres at both ends, 8 chromosomes had telomeres at one end, and only one chromosome lacked telomeric sequences at either end. Of the two unplaced short contigs, one (4MBase) is suggested by sequence similarity to be a fragment of centromeric satellite sequence, and the other (0.4MBase) an rDNA fragment. Reads from a *C. rubella* accession (SRR8394204) aligned to the genome with 99.0% completeness and reads from a *C. grandiflora* accession (ERR14012352) aligned with 92.9% completeness. Similarly, an ascension of *C. orientalis* (SRR6179227) aligned with 98.3% completeness. These sequence alignments showed an increase in *C. orientalis* mapping at the end of Chromosome 10 of our assembly and an increase in *C. rubella* and *C. grandiflora* mapping at the end of Chromosome 2; evidence of homeologous exchange between the two chromosomes (Supplemental Figure 2).

### Genetic structure at the collection site level in NYC

We used a genomic principal component analysis (PCA) and population genetic statistics to assess differentiation between sampling sites in NYC. In the PCA, individuals within sampling sites are tightly clustered with one another (Figure 1A). Population structure in principal component (PC) space is unchanged when evaluating the subgenomes separately (Supplemental Figure 3). Total nucleotide diversity (π) within sampling sites ranged from 0.000155 to 0.0379 on the *C. grandiflora*-like subgenome and 0.000144 to 0.0235 on the *C. orientalis*-like subgenome [Table S3]. Some sampling sites had higher nucleotide diversity than others and this appeared to be due to both higher individual heterozygosity and reduced relatedness among samples within the site (Supplemental Figures 4-7). The first PC separates the Newark, New Jersey site from the rest of the sites within the state of New York, while the second and third principal components show variation within the boroughs (Figure 1A and Figure 1B). There does not appear to be significant clustering by borough among the New York state samples (Supplemental Figure 8), though samples from sites 14 (NY_09 and NY_10), 4 (NY_45, NY_46, NY_47), and 21 (NY_32, NY_33, NY_34) are disjointed from the main cloud of samples. Among the NYC sites, there is moderate genetic differentiation; Fst between all sampling sites is -0.06213 to 0.91501 with the highest values of differentiation being driven by sites 24 (Newark, NJ), 21, and 14 [Table S4]. Across the NYC metropolitan area, we did not find evidence of increasing genetic distance with geographic distance (Figure 1D; mantel test statistic, r = 0.117; p-value = 0.275). Among the NYC *C. bursa-pastoris*, average nucleotide diversity (π) was 0.02206 at 0-fold sites (0.01825 *C. grandiflora* subgenome; 0.02554 C. orientalis subgenome) and 0.0490 at 4-fold degenerate sites (0.03588 C. grandiflora subgenome; 0.06044 *C. orientalis* subgenome).

**Figure 1:**
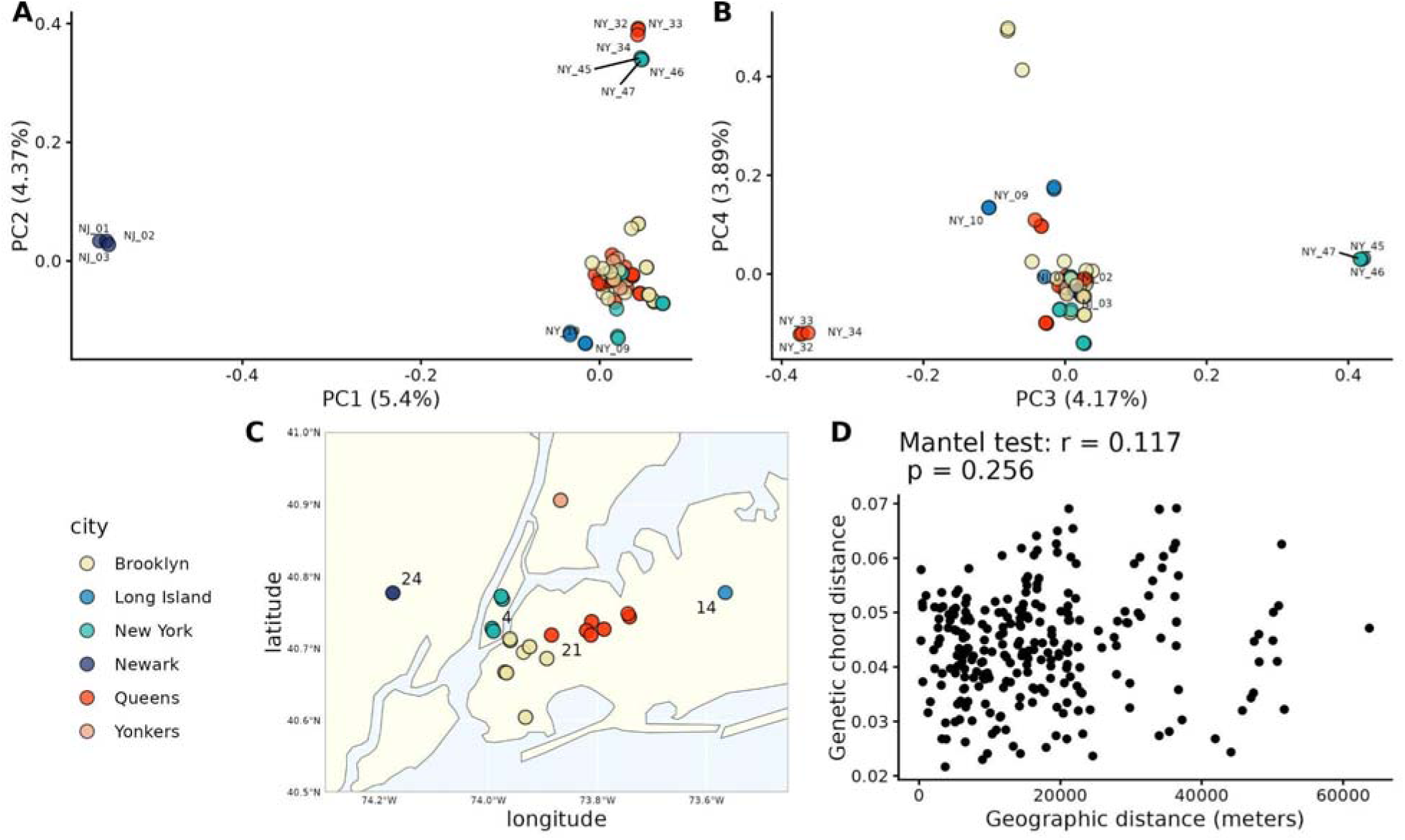
Genetic structure of *Capsella bursa-pastoris* in the New York Metropolitan area. Colors indicate cities within NYC. **A:** First and second principal component of all *C. bursa-pastoris* in NYC. Individuals from sampling sites 4 (NY_45, NY_46, NY_47), 14 (NY_09 and NY_10), 21 (NY_32, NY_33, NY_34), and 24 (NJ_01, NJ_02, NJ_03) are labelled. **B:** Third and fourth principal components of *C. bursa-pastoris*. **C:** Map of sampling locations in the NYC metro area. **D:** Geographic distance and genetic distances between pairs of sampling locations. Alt text: Multipanel figure showing genetic population structure of *Capsella bursa-pastoris* in the New York City metropolitan area. Panels A and B are colorful scatterplots, Panel C indicates individuals with colorful points on a map, and Panel D is a colorless scatterplot annotated with the results of a Mantel test showing no statistically significant relationship between geographic distance and genetic chord distance.

### NYC *C. bursa-pastoris* is most closely related to northern Eurasian *C. bursa-pastoris*

In our principal component analysis, we find that *C. bursa-pastoris* from NYC clusters most closely with *C. bursa-pastoris* from northern European locations (Figure 2B, 2C). Mean Fst between NYC and Northern Eurasia is 0.132 (0.1241 C. grandiflora subgenome; 0.1398 *C. orientalis* subgenome) while mean Fst comparing NYC the Mediterranean and East Asian groups are 0.522 (0.5010 *C. grandiflora* subgenome; 0.54403 *C. orientalis* subgenome) and 0.686 (0.6529 *C. grandiflora* subgenome; 0.7177 *C. orientalis* subgenome), respectively [Table S5]. PC1 separates *C. bursa-pastoris* geographically where NYC and Northern Eurasian *C. bursa-pastoris*, Mediterranean *C. bursa-pastoris*, and East Asian *C. bursa-pastoris* form groups while a single sample from Kyrgyzstan is intermediate between the three groups. PC3 further differentiates individuals in the Northern Eurasian group from one another (Figure 2B, 2C) with NYC *C. bursa-pastoris* forming a distinct group. Principal component analysis performed with only members of the Northern Eurasian cluster reveals further structure within Northern Eurasia and highlights differences between Northern Eurasian *C. bursa-pastoris* and NYC *C. bursa-pastoris* (Supplemental Figure 9). ADMIXTURE among the Northern Eurasian groups with downsampled NYC further support the separation of NYC from other Northern Eurasian samples and indicate New Jersey as being more closely related than New York samples are to ancestral Northern Eurasian genotypes (Supplemental Figure 10).

**Figure 2:**
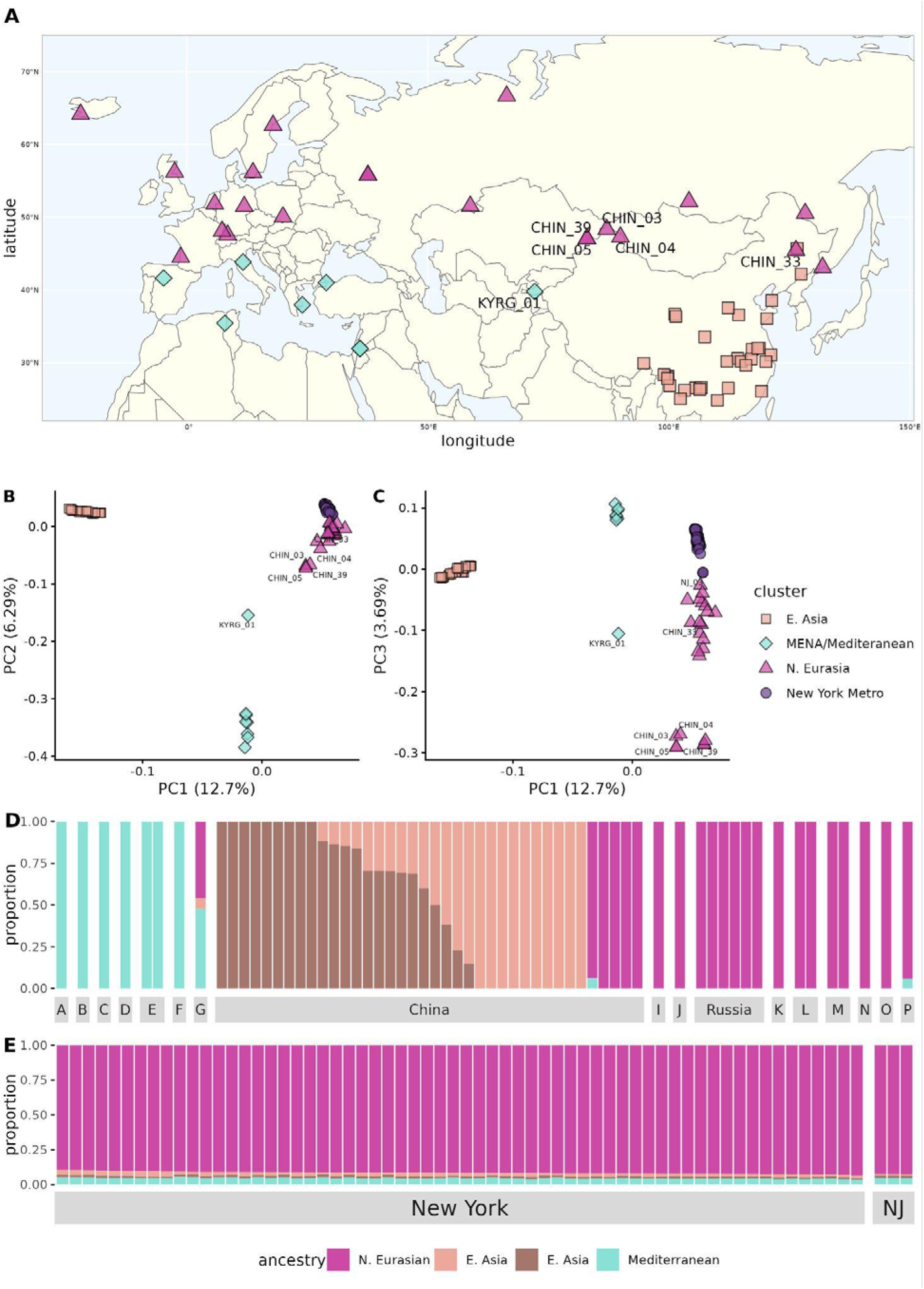
Population structure of *Capsella bursa-pastoris* in ancestral range. **A:** Map of whole genome sequences from the ancestral range. **B:** First and second principal components of NYC *C. bursa-pastoris* and ancestral range *C. bursa-pastoris*. **C:** First and third principal components of NYC *C. bursa-pastoris* and ancestral range *C. bursa-pastoris*. **D**: Ancestry proportions of *C. bursa-pastoris* in ancestral range given by ADMIXTURE when K=4. Individuals are colored by the proportion of ancestry. Facet labels indicate country (A=Algeria, B=Spain, C=Greece, D=Italy, E=Jordan, F=Turkey, G=Kyrgyzstan, H=Taiwan, I=Switzerland, J=Poland, K=Germany, L=Sweden, M=France, N=Scotland, O=Netherlands, P=Iceland) **E:** Ancestry proportions of NYC where ancestry is defined by allele frequencies within populations from the ancestral range. Alt text: Multipanel figure showing the relationship between *Capsella bursa-pastoris* in the New York City metropolitan area and *Capsella bursa-pastoris* in the ancestral range across Eurasia and north Africa.

Our ADMIXTURE analysis of *C. bursa-pastoris* in the ancestral range showed K=4 groups had the lowest cross validation error rate (Supplemental Figure 11). Members of the four groups in the ADMIXTURE analysis correspond to PCA clustering insofar as the N. Eurasian, Mediterranean, and East Asian samples are grouped, with Kyrgyzstan showing the most intermediate ancestry between all three (Figure 2D). The ADMIXTURE analysis further separated the East. Asian population into two groups (Figure 2D). Samples from central Asia (northwestern China and Kyrgyzstan) as well as some European genotypes (ICEL_01 from Iceland and FRAN_02 from France) all show mixed ancestry proportions across the three major groups (Figure 2D). When NYC individuals are assigned population proportions based on the ancestral groups, all NYC *C. bursa-pastoris* show a majority ancestry proportion from the Northern Eurasian group with moderate ancestry from the Mediterranean and East Asian group (Figure 2E).

### Evaluation of introgressed regions in NYC *C. bursa-pastoris*

We evaluated NYC *C. bursa-pastoris* for past introgression from *C. rubella* and identified specific regions introgressed from *C. rubella* in the *C. grandiflora*-like subgenome. We find NYC *C. bursa-pastoris* has genomic regions inferred to be introgressed from *C. rubella* (Figure 3B and Supplemental Figure 12). Within the population, introgressed regions vary in length and frequency (Figure 3B) and, on average, 14% of the genome is *C. rubella* ancestry [Table S6]. The region of Chromosome 10 that was indicated to be derived from the *C. orientalis* subgenome due to homeologous exchange was not inferred to have *C. rubella* ancestry (Figure 3B). East Asian *C. bursa-pastoris* that were evaluated for introgression as a control had no *C. rubella* ancestry inferred (Supplemental Figure 13). The principal component analysis of putatively introgressed SNPs showed that NYC *C. bursa-pastoris* individuals were more similar to *C. rubella* individuals than *C. orientalis*, *C. grandiflora*, or East Asian *C. bursa-pastoris* (Supplemental Figure 14). At non-introgressed SNPs, NYC C. bursa-pastoris clustered with East Asian *C. bursa-pastoris* samples (Supplemental Figure 15). We compared the proportion of basepairs from genes within introgressed regions to basepairs from genes across the *C. grandiflora*-like subgenome where there was sufficient mappability. We found that introgressed regions contained more basepairs from genes than expected by chance (Figure 3C). Lastly, nucleotide diversity is higher at SNPs that are putatively introgressed from *C. rubella* than at SNPs that are not introgressed; average pi is 0.125 at introgressed SNPs compared to 0.08 at non-introgressed SNPs (Figure 3D; p-value = 6.82e^-12^). Additionally, nucleotide diversity per SNP is higher among individuals that are introgressed at a SNP that individuals who are not; average pi across introgressed SNPs is 0.073 for introgressed individuals and 0.013 for non-introgressed individuals (Figure 3E; p-value = 4.55e^-99^).

**Figure 3:**
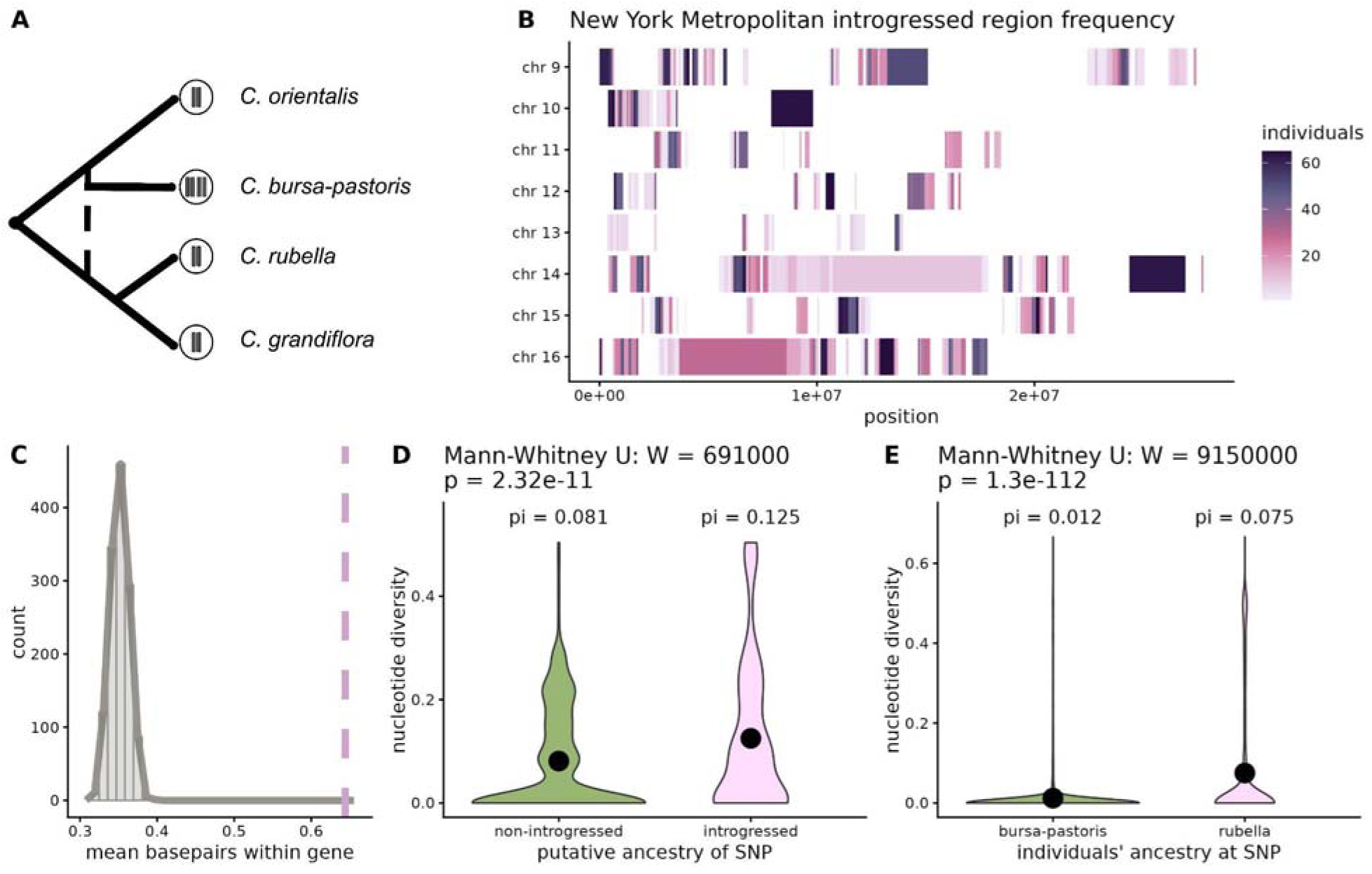
Introgression in the NYC metropolitan area. **A:** schematic species tree for the *Capsella* genus. **B:** Introgressed regions across chromosomes of NYC metro *C. bursa-pastoris*. Shading indicates the number of NYC individuals sharing an introgressed region where darker color indicates more individuals share an introgressed region. **C:** The mean proportion of basepairs in genes within a sample of genomic regions. A distribution of the mean proportion of genic basepairs from random samples of the genome, where sampled regions are the same size as introgressed regions (in grey). Dashed vertical line indicates the mean proportion of genic basepairs within introgressed regions. **D**: Nucleotide diversity across all individuals at putatively introgressed SNPs (right; pink) and putatively non introgressed SNPs (left; green). Black dots show average value within a group. **E**: Nucleotide diversity among introgressed (right; pink) and non introgressed (left; green) individuals at evaluated SNPs. Black dots show average value within a group. Alt text: Detailed graphs showing introgression in *Capsella bursa-pastoris* from the New York metropolitan area. Panels D and E are annotated with average nucleotide diversity values for each group.

## Discussion

Here, we explore the genetic diversity of *C. bursa-pastoris* in the New York metropolitan area and present a reference genome for the species to gain insight to the evolutionary forces that shape variation in this urban environment to understand how introgression influences genetic variation in the introduced range.

We generated a chromosome-level whole genome assembly for *C. bursa-pastoris* from a single individual collected in Yonkers, New York. This genome represents haplotypes within a temperate region the introduced range on the East Coast of the United States of America. Importantly, another chromosome-level assembly for this species is published and represents an individual collected in part of the ancestral range of the species, Moscow, Russia (Penin et al. 2024). With the addition of our assembly, these resources provide an exciting opportunity to capture a larger amount of genetic variation. The aligned sequencing depth of *C. rubella*, *C. grandiflora*, and *C. orientalis* revealed a balanced homeologous exchange between Chromosome 2 and Chromosome 10 of our assembly (Supplemental Figure 2). The high degree of synteny between our assembly and other *C. bursa-pastoris* assemblies (Fiscus 2022; Penin et. al. 2024) (Supplemental Figure 1) suggests this genomic feature is widespread in the species. Our results highlight that *C. bursa-pastoris* is a highly diverse species where the NYC population is clearly diverged from even its closest relatives. Therefore, sequencing efforts across the broad geographic range of *C. bursa-pastoris* may aid the construction of a pangeome which allows this species to become a useful resource in the field of population genetics (Roberts et al. 2025).

To investigate population structure in a region of the introduced range, we used principal components to analyze the whole genome sequences of *C. bursa-pastoris* within the NYC metropolitan area (NYC). We find that within NYC, individuals are most similar to those in their sampling location. Thus, patches are likely colonized by single individuals that then proliferate via self-fertilization, similar to urban *Arabidopsis thaliana* (Schmitz et al. 2024). Individuals from the state of New York are more closely related to one another than they are to individuals at the sampled location in Newark, New Jersey; mean Fst between NJ and NY populations is 0.497 to 0.932 [Table S4]. This might suggest that, for small urban plants, geographic barriers, such as the Hudson River, provide a reduction in gene flow despite anthropogenic influence. Nevertheless, the PCs explain very little variation in genotypes; cumulative variance explained by the first 10 PCs is 38.21%. Therefore, despite being able to detect some differentiation between NY and NJ genotypes in a PCA, we do not find evidence of strong population structure broadly across the New York City Metropolitan area. In agreement with these findings, we do not detect a signal of isolation-by-distance among sampling locations within NYC. Isolation-by-distance is the expected result of a stepwise pattern for colonization assuming individual dispersal distances and genetic drift are sufficiently strong forces structuring populations. Previous studies of *C. bursa-pastoris* in the ancestral range also do not detect a pattern of isolation-by-distance within the ancestral groups (Cornille et al. 2016). In urban populations generally, it is not uncommon to fail to detect a pattern of isolation-by-distance (Tsutsui and Case 2001; Platt et al. 2010; Guo et al. 2018). It appears that other forces, likely in conjunction with human-mediated dispersal, primarily drive the genetic structure of *C. bursa-pastoris* in NYC.

To contextualize the diversity of NYC *C. bursa-pastoris*, we compared our samples from the NYC metropolitan area to *C. bursa-pastoris* in its ancestral range. We find that NYC *C. bursa-pastoris* is closely related to *C. bursa-pastoris* from Northern Eurasia. The mean genome-wide Fst between NYC and Northern Eurasia is 0.132 (mean Fst is 0.1241 on the *C. grandiflora*-like subgenome 0.1389 on the *C. orientalis*-like subgenome [Table S5]). This result is supported by previous studies of the *C. bursa-pastoris* which conclude that global *C. bursa-pastoris* genetic populations have an association with regional climate (Wesse et al 2021, Neuffer and Hurka 1999). Temperate New York and New Jersey would therefore be most closely related to other temperate *C. bursa-pastoris* populations in the ancestral range. Additionally, these results are in concordance with the history of European colonization and immigration on the east coast of the United States. Previous studies posited that *C. bursa-pastoris* populations in North America were introduced without much change (Neuffer and Hurka 1999; Wesse et al. 2021). However, despite the close association, our analyses show the NYC population of *C. bursa-pastoris* noticeably deviates from the N. Eurasian cluster in a PCA, making NYC *C. bursa-pastoris* appear to be a distinct population when compared to its closest sampled relatives. This is in contrast to similar investigations of introduced populations in cities, where individuals from a city colocalize in principle component analyses with their putative source populations (Combs et al. 2018; Schmitz et al. 2024). This apparent divergence from populations in the ancestral range could be due to the introduction of many source lineages with subsequent hybridization in NYC. Though identifying the sources for populations in cosmopolitan species is known to be difficult (Estoup and Guillemaud 2010; Cristescu 2016), species introductions to new ranges may influence the rate of hybridization by eroding environment-independent pre-zygostic barriers to mating (Vallejo-Marín and Hiscock 2016). Alternatively, there remains the possibility that lineages in the ancestral range are undersampled in comparison to the NYC group. As is the case for human populations in Newfoundland and Labrador (Zhai et al. 2016), diverse sampling may provide useful data for contextualizing differences between the N. Eurasian and NYC groups. Nevertheless, our data proved additional evidence that introductions of this species caused populations in the introduced range to evolve divergently from those in the ancestral range.

Finally, we explored the role of past interspecific introgression on the establishment of *C. bursa-pastoris* in the introduced range. There is extensive evidence of introgression from *C. rubella* to *C. bursa-pastoris* in its ancestral range of Eurasia (Slotte et al. 2008; Han et al. 2015; Kryvokhyzha et al. 2019) where introgression is restricted to populations in northern and western Eurasia. As with other extant populations in northwestern Eurasia, we find NYC *C. bursa-pastoris* has evidence of introgression from *C. rubella*. Previous studies investigating introgression were based on reduced representation sequencing and/or aligned *C. bursa-pastoris* samples to a *C. rubella* reference genome. With an excess of densely sampled, high quality genetic data, we are able to explore genetic variation in NYC *C. bursa-pastoris* on a finer scale and potentially capture cryptic introgression that is difficult to identify through other means, such as limited genetic data, morphology, or cytology (Brown et al. 2024). Additionally, our data allow us to determine with precision which genomic regions are introgressed and evaluate particular loci in future studies to understand their maintenance in this population. Introgressed genomic regions vary in rarity and size in NYC which indicates a complex history of introgression in the ancestral range.

Given the similarity between *C. rubella* and *C. grandiflora*, it is difficult to rule out the possibility that the introgression detected in this study is definitively from *C. rubella*. The founding haplotypes of *C. rubella* appear to be a random sampling of the diversity of *C. grandiflora* (Brandvain et al. 2013). While we cannot completely rule out the possibility that the introgression we detect is from *C. grandiflora*, there are several lines of evidence that point to the source of introgressed material being *C. rubella*. Tests of incomplete lineage sorting show *C. rubella* has higher likelihoods of gene flow with *C. bursa-pastoris* than does *C. grandiflora* (Kryvokhyzha et al. 2019). Additionally, haplotype networks of all four species in the *Capsella* genus show haplotype sharing among *C. rubella* and *C. bursa-pastoris* that is not present in *C. grandiflora* (Han et al. 2015). In artificial crosses, *C. bursa-pastoris* and *C. grandiflora* do not produce viable seeds (Kryvokhyzha et al. 2019). Moreover, there is evidence of fairly strong reproductive isolation between the diploid species (Han et al. 2015; Sicard et al. 2015; Lafon-Placette et al. 2018; Kryvokhyzha et al. 2019; Dziasek et al. 2024). Nonetheless, *C. grandiflora* and *C. orientalis* can produce viable offspring that spontaneously become allotetraploids (Bachmann et al. 2021) suggesting that mating between these two diploid species could have contributed to the formation of *C. bursa-pastoris*. While *C. rubella* and *C. grandiflora* co-occur in Greece, *C. orientalis* does not have a known overlapping range with either of the other diploids. Therefore, it is unlikely that gene flow between *C. orientalis* and *C. grandiflora* or *C. rubella* could produce the pattern we see here in our data.

Due to the limited geographic range of *C. rubella*, we can confidently restrict the introgression event(s) to the ancestral range prior to this species introduction to the United States. Therefore, we asked what role the introgressed genomic regions play in evolution within the introduced range. An intuitive explanation is that introgressed genetic material is maintained because it is beneficial in the population. Alternatively, introgressed material could be neutral or weakly deleterious but remain within the population due to insufficient time for recombination and natural selection to remove it (Moran et al. 2021; Aguillon et al. 2022). In this second scenario, we might expect stronger evidence of introgression in regions depleted of genes. Instead, we find that introgressed regions of the genome contain more basepairs derived from genes than similarly sized regions in the genome. Though a thorough investigation of this scenario is beyond the scope of this paper, introgressed regions in NYC *C. bursa-pastoris* show a high degree of genetic variation which suggest introgression in the past was widespread and/or occurred in multiple pulses. In addition, we find that introgression increases genetic variation in general. Past introgression from *C. rubella* might have played a role in the establishment of *C. bursa-pastoris* in the introduced range by providing additional genetic variation beyond the ancestral *C. bursa-pastoris* haplotypes. We find that introgression from *C. rubella* contributes to nucleotide diversity in *C. bursa-pastoris* not only by introducing alternative alleles at loci but the introgressed regions themselves are also genetically diverse.

## Materials and Methods

### *Capsella bursa-pastoris* genome assembly and annotation

Genomic DNA was extracted from the leaf tissue of a single individual of *Capsella bursa-pastoris* from New York (Latitude: 40.906082, Longitude: -73.867637; *Yonkers, New York*) and sequenced at the University of Delaware using the Pacific Biosciences HiFi platform. We generated the assembly using HiFisam (version 0.19.8) (Cheng et al. 2021) with the parameter - k 61. Gene-space completeness was determined at the genomic level using the Brassicales_odb12 database from BUSCO (v.5, (Manni et al. 2021). We used COGE Synmap (Lyons et al. 2008; Albert and Krabbenhoft 2023) and D-Genies (Cabanettes and Klopp 2018) to compare our assembly to other *C. bursa-pastoris* assemblies (Fiscus 2022). To further investigate intergenic regions and identify the subgenomes, we aligned genomic DNA from *C. rubella*, *C. grandiflora,* and *C. orientalis* (*C. rubella* SRR8394204; *C. grandiflora* ERR14012352; *C. orientalis* SRR6179227) with BWA mem (Li 2013). Genome annotation was performed with AUGUSTUS (Stanke et al. 2006) using *Arabidopsis thaliana* as the species database. We identified 0-fold and 4-fold sites with Degenotate (Mirchandani et al. 2024).

### Population sampling and DNA sequencing

*C. bursa-pastoris* maternal lines were collected around the New York Metropolitan area in 2017 (Figure 1C). Maternal lines were maintained by R. Panko and allowed to self-fertilize for 2-3 generations. For this study, we germinated seeds from these lines on a soil mix (2 SureMix:1 RediEarth) in growth chambers at Michigan State University (East Lansing, Michigan). Young leaf tissue was collected from plants at the rosette stage and frozen at -80. Genomic DNA was extracted using the Qiagen DNEasy Plant Mini kit (Germantown, Maryland). Extracted DNA was sequenced at Michigan State University Genomics Core. Libraries were prepared using the Illumina TruSeq Nano DNA Library Preparation Kit with IDT for Illumina Unique Dual Index adapters following manufacturer’s recommendations. Completed libraries were assessed for quality and quantified using a combination of Qubit dsDNA HS and Agilent 4200 TapeStation HS DNA1000 assays. The libraries were pooled in equimolar amounts and this pool was quantified using the Invitrogen Collibri Quantification qPCR kit. The pool was loaded onto one lane of an Illumina NovaSeq 6000 S4 flow cell and sequencing was performed in a 2x150bp paired end format using a standard v1.5 NovaSeq reagent cartridge. Base calling was done by Illumina Real Time Analysis (RTA) (v3.4.4) and the output of RTA was demultiplexed and converted to FastQ format with Illumina Bcl2fastq (v2.20.0). In total, we sequenced 65 individuals from the New York metropolitan area (herein ‘NYC’) to an average read length of 150 bp and 30x coverage. These individuals represented 23 sampling locations around New York and New Jersey, USA [Table S1].

### Alignment, variant calling and filtering

We generated two VCF files, a single-species VCF and multi-species VCF, for our analyses. The single-species VCF contained data from *C. bursa-pastoris* samples which we obtained by combining the 65 NYC samples with 68 previously sequenced and publicly available whole genome sequences of *C. bursa-pastoris* [Table S2]. With raw fastq files representing 133 *C. bursa-pastoris* samples, we trimmed adapters and filtered reads for quality. We aligned the quality filtered reads to our genome assembly with bwa-mem (Li 2013). After alignment, we called variant sites and maintained invariant sites using GATK (McKenna et al. 2010; Van der Auwera and O’Connor 2020) for downstream population genetic analyses.

We filtered the single-species VCF, containing only *C. bursa-pastoris* individuals, following GATK recommendations for hard quality filtering (Van der Auwera and O’Connor 2020). We used BCFtools (version 1.18) to remove variants with a quality-by-depth ratio less than 2 (QD < 2), PhredScaled probability of strand bias above 60 (FS > 60), strand odds ratio greater than 3 (SOR > 3), RMS mapping quality less than 40 (MQ < 40), mapping quality rank sum less than - 12.5 (MQRankSum < -12.5), and read position rank sum less than -8.0 (ReadPosRankSum < - 8.0). To identify multicopy genomic regions, we used ParaMask (Tjeng et al. 2025) on biallelic SNP variant sites. Then, we removed sites with a frequency of missing calls over 10%, quality scores that were missing or below 20, and more than two alleles present. Further, we removed SNPs identified by ParaMask to be in multicopy regions and filtered for depth; we removed sites with a read depth value above or below 1.5 times the interquartile range (below 3753.5 and above 6133.5). We then separated invariant and variant sites to remove variant sites with a minor allele frequency less than 0.025 and insertion/deletions (indels), before recombining the invariant and variant sites. After filtering, we were left with 1,457,611 biallelic SNPs and 5,023,509 invariant sites, where invariant sites were monoallelic SNPs or sites without known alternate alleles. This single-species VCF was used for all population genetic analyses of *C. bursa-pastoris*.

The multispecies VCF contained samples from the four major species in the *Capsella* genus: *C. bursa-pastoris*, *C. rubella*, *C. grandiflora*, and *C. orientalis* (Figure 3A). As above, we performed quality filtering on the raw fastq files and aligned them to our reference assembly with bwa-mem. In order to mitigate the impact of mismapped reads, we aligned the species to our assembly in the following manner: First, we aligned 133 *C. bursa-pastoris* individuals to the entire *C. bursa-pastoris* reference genome so that reads from this allotetraploid species may align to either subgenome based on best mapping location. DNA libraries from the diploid species were aligned to only the *C. grandiflora*-like subgenome of our reference (Scaffolds 9-16). We aligned 50 *C. rubella* and 68 *C. grandiflora* samples to our assembly and called variants in *C. bursa-pastoris*, *C. rubella*, and *C. grandiflora* jointly. After filtering for quality (described below), we retained 5,270,306 SNPs and used these data to identify introgression in *C. bursa-pastoris*. We called variants for 31 *C. orientalis* samples separately and maintained invariant sites before intersecting the dataset with SNPs across the other three species. We did this to reduce the risk of miscalling variants due to the potential poor alignment of *C. orientalis* to the *C. grandiflora*-like subgenome and to capture sites that may be polymorphic across *C. bursa-pastoris*, *C. rubella*, and *C. grandiflora* but monomorphic in *C. orientalis*. We used this final multi-species VCF (897,650 SNPs) to validate the origin of introgression in *C. bursa-pastoris*.

For the multispecies VCF, we completed the same variant filtering steps as in the single species VCF with minor alternations. In brief, we removed sites that did not pass the GATK Best Practice hard filters (noted above), contained more than 10% missing calls, and had a read depth below the minimum or above the maximum depth threshold. We excluded *C. grandiflora* samples from site level heterozygosity calculations as *C. grandiflora* is an obligate outcrosser with elevated heterozygosity. Among the remaining genotypes from selfing species (*C. bursa-pastoris, C. rubella*, and *C. orientalis*), we removed 2 individuals with unusually high heterozygosity. Lastly, we filtered sites to keep only those with a within-species minor allele frequency above 0.025 in the selfing species (*C. bursa-pastoris*, *C. rubella*, and *C. orientalis*) and above 0.05 in *C. grandiflora*.

### Population structure of *C. bursa-pastoris* in NYC

We used principal component analysis (PCA) to analyze the population structure of *Capsella bursa-pastoris* within NYC. We removed sites in linkage disequilibrium (sites with an r^2^ correlation above 50% in windows of 500 SNPs and a step size of 50 SNPs) in NYC *C. bursa-pastoris* and returned all available principal components (n=65) to accurately capture the variance explained by each. Linkage disequilibrium calculations, site pruning, and principle component vector analyses were performed with PLINK (v2.00 (Chang et al. 2015)) using only variant sites in the single-species VCF. We performed the analysis twice: once with all samples in the NYC metropolitan area and once excluding our samples from New Jersey from the linkage disequilibrium calculations and the PCA. We extracted all available principal components (n=62) for the PCA that excluded New Jersey samples.

To assess genetic diversity in the NYC population, we calculated Hudson’s Fst (Hudson 1992; Bhatia et. al. 2013) between sampling sites in the NYC metropolitan area (n=24 sampling sites; 2-3 individuals per sampling site) using PLINK. We then calculated nucleotide diversity (π) among the NYC metropolitan area *C. bursa-pastoris* in non-overlapping windows of 10 thousand basepairs each along the genome using PIXY (v2.0.0.beta14) (Korunes and Samuk 2021). We assessed nucleotide diversity at 4-fold sites, 0-fold sites, and all sites across the whole genome and per subgenome. We calculated the genome-wide average of nucleotide diversity in the NYC metro population as the sum of differences divided by the sum of comparisons across all windows.

### Test for Isolation-by-distance patterns in NYC *C. bursa-pastoris*

We tested for a pattern of isolation-by-distance in the NYC populations in R (v4.5.1; (R Core Team 2025)). Using genotypes at 4-fold degenerate sites, we calculated Cavalli-Sforza and Edwards’ genetic chord distance between pairs of sampling sites within NYC using the R package *hierfstat* (v.0.5-11; (Goudet 2005)). We chose to calculate genetic chord distance as it has the most power to detect isolation-by-distance (Séré et al. 2017). Geographic distances were calculated from sample latitude and longitude using the *sf* package (Pebesma 2018; Pebesma and Bivand 2023). We found the correlation between the genetic and geographic dissimilarity matrices by calculating the Mantel statistic with the package *vegan* (v.2.7-2; (Oksanen et al. 2025).

### Relationship of NYC *C. bursa-pastoris* to ancestral range *C. bursa-pastoris*

Similar to the analysis within NYC, we assessed the population structure of *C. bursa-pastoris* with a PCA. Again, we used PLINK to prune sites in linkage disequilibrium and calculate principal components, this time combining the NYC individuals with those from the ancestral range (n=133 individuals). We then evaluated individual ancestry proportions among *C. bursa-pastoris* from the ancestral range with ADMIXTURE (v1.3.0 (Alexander et al. 2009) allowing K-groups to vary between 1 and 6. To contextualize NYC *C. bursa-pastoris* in the ancestral range, we used the allele frequencies from populations in the ancestral range to assign ancestry to the NYC *C. bursa-pastoris* population.

Based on the results of the PCA of all *C. bursa-pastoris* individuals, we performed an additional PCA and ADMIXTURE analysis with only 24 samples from NYC (1 individual per sampling site) and 24 samples from Northern Eurasia. We pruned sites in linkage disequilibrium separately for this subset of samples and ran ADMIXTURE allowing K-groups to vary from 2-10.

### Local ancestry inference

Our reference assembly and surplus of whole genome sequencing data at sufficient depth allow for increased granularity in our investigation of NYC *C. bursa-pastoris* ancestry. Therefore, to define local ancestry along chromosomes of *C. bursa-pastoris*, we used Ancestry HMM (Corbett-Detig and Nielsen 2017). Ancestry HMM compares allele frequencies from a query population to the allele frequencies in user-defined parental populations. Given prior research in *C. bursa-pastoris* (Kryvokhyzha et. al. 2019), we looked for evidence of introgression from *C. rubella* into *C. bursa-pastoris* on the *C. grandiflora*-like subgenome (scaffold 9-16 of our assembly). We aimed to distinguish *C. rubella* ancestry from *C. bursa-pastoris* ancestry in the population. Therefore, our parental populations were defined as 50 *C. rubella* genotypes, along with 30 *C. bursa-pastoris* genotypes from East Asia. We selected East Asian samples based on our population genetic analyses and research from other groups that suggests East Asian *C. bursa-pastoris* has not received any introgressed material from the *C. rubella* or *C. grandiflora* lineage (Huang et al. 2012; Han et al. 2015; Kryvokhyzha et al. 2019). Though our subgenome of interest is derived from a *C. grandiflora*-like ancestor (Douglas et al. 2015), we did not choose *C. grandiflora* as a parent because modern day *Capsella grandiflora* is likely to have diverged from those that contributed to the formation of *C. bursa-pastoris*.

From our quality filtered multispecies VCF, we identified SNPs in linkage disequilibrium within the parental populations. We used PLINK2 with the same parameters in analysis of population structure to calculate linkage disequilibrium (an r^2^ correlation above 50% in windows of 500 SNPs and a step size of 50 SNPs) and removed those SNPs with BCFtools. We calculated linkage disequilibrium within the parental populations separately because we are interested in capturing the signal of SNPs in linkage disequilibrium as a result of introgression, as opposed to those that are in linkage disequilibrium because they are linked in the parental populations. We ran Ancestry HMM with the following parameters: 2 ancestral populations with admixture proportions of 0.3 and 0.7 (estimated from (Han et al. 2015)), a pulse of *C. rubella* ancestry 50,000 generations ago, while estimating the proportion of present ancestry from *C. rubella* in the modern day population.

### Evaluation of *Capsella rubella* ancestry

We evaluated the signal of *C. rubella* ancestry in two ways. First, we included 5 *C. bursa-pastoris* individuals from East Asia in the population to be evaluated for *C. rubella* ancestry. We expect no *C. rubella* ancestry in these individuals based on prior research (Huang et al. 2012; Han et al. 2015; Kryvokhyzha et al. 2019) and because we are using the East Asian population to define *C. bursa-pastoris* ancestry. Second, we performed principal component analyses at *C. rubella* introgressed SNPs and non-introgressed SNPs. In the PCA*, C. bursa-pastoris* individuals should cluster more closely with *C. rubella* individuals in principal component space at SNPs identified to be introgressed.

### Introgressed region content and diversity

We examined the content of introgressed regions with analyses of genic content and nucleotide diversity. To estimate genic content, we randomly sampled from regions of the genome that are the same size as introgressed regions and calculated the mean number of basepairs that fell within a gene. We restricted our random sampling of the genome to mappable regions by excluding regions with zero SNPs in a 10kb window. We repeated this sampling 1326 times to generate a null distribution. We then compared the mean proportion of gene base pairs in introgressed regions to our null. We used BEDtools *annotate* (v2.31) to determine the proportion of each region (random or introgressed) that contained basepairs originating from genes.

To determine if introgression contributed to genetic diversity in NYC *C. bursa-pastoris*, we evaluated nucleotide diversity in two ways. Firstly, we compared nucleotide diversity at putatively introgressed SNPs and putatively non-introgressed SNPs. We defined a SNP as putatively introgressed if more than 80% of NYC individuals had a posterior probability of *C. rubella* ancestry above 0.5 in our local ancestry analysis. Putatively non-introgressed SNPs were those where more than 80% of individuals had a posterior probability of *C. bursa-pastoris* ancestry above 0.5. Secondly, at all ancestry evaluated SNPs, we calculated nucleotide diversity among NYC individuals who were introgressed at that locus versus those not introgressed at the same locus. In both analyses, we determined significance between the groups using a Mann-Whitney U test implemented in R.

## Data Availability

The genomic data in this article are available from the NCBI Short-Read Archive at [URL], and can be accessed with [Bioproject Number]. The data and analysis scripts underlying this article are available in a GitHub Repository archived on Zenodo, available at [DOI].

## Acknowledgments

These authors thank members of the Josephs Lab and David Lowry for helpful comments. Research reported was supported by the National Institute of General Medical Sciences of the National Institute of Health (NIH) under award R35GM142829 to EBJ and by the National Science Foundation Graduate Research Fellowship under DGE-1848739 to MKWB. This work was supported in part through computational resources and services provided by the Institute for Cyber-Enabled Research at Michigan State University.

## Notes

https://github.com/mwilsonbrown/cbp_nyc_variation

### Competing Interest Statement

The authors have declared no competing interest.

### Summary of Updates

Minor revisions to text; additional significant statement added

