## Supplemental Figures for "Introgression across ploidies contributes to genetic diversity in introduced urban *Capsella bursa-pastoris*"

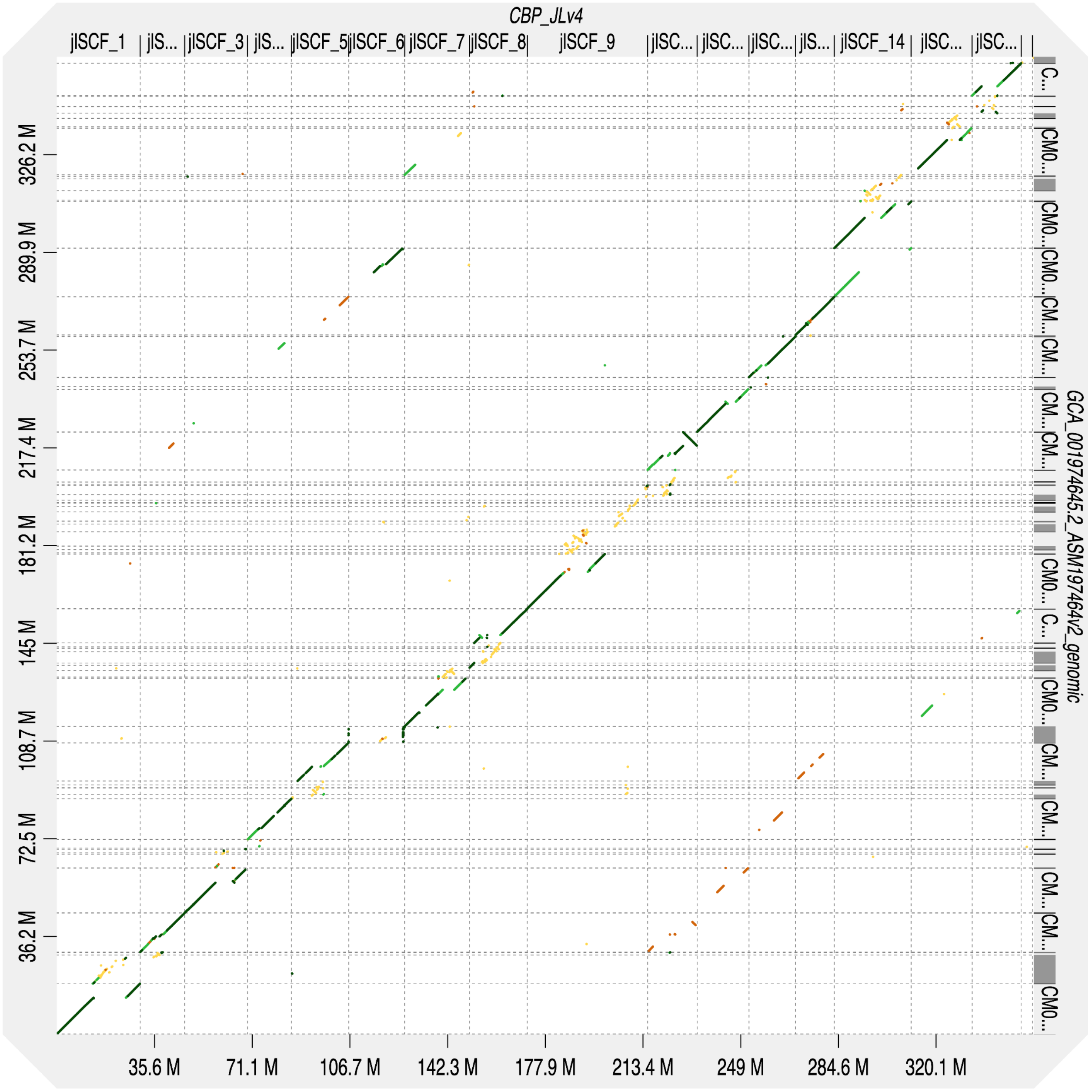


Supplemental figure 1: Synteny between assembly referenced in manuscript (vertical axis) and generated in Penin et. al. 2024 (horizontal axis) generated with D-Genies. D-Genies (WGS-v-WGS, Id>=0.4), CBP_JL4 v ASM197464v2; Josephs Lab, Lomonosov Moscow State University

Alt text: Dot plot showing similarity between two genome assemblies.

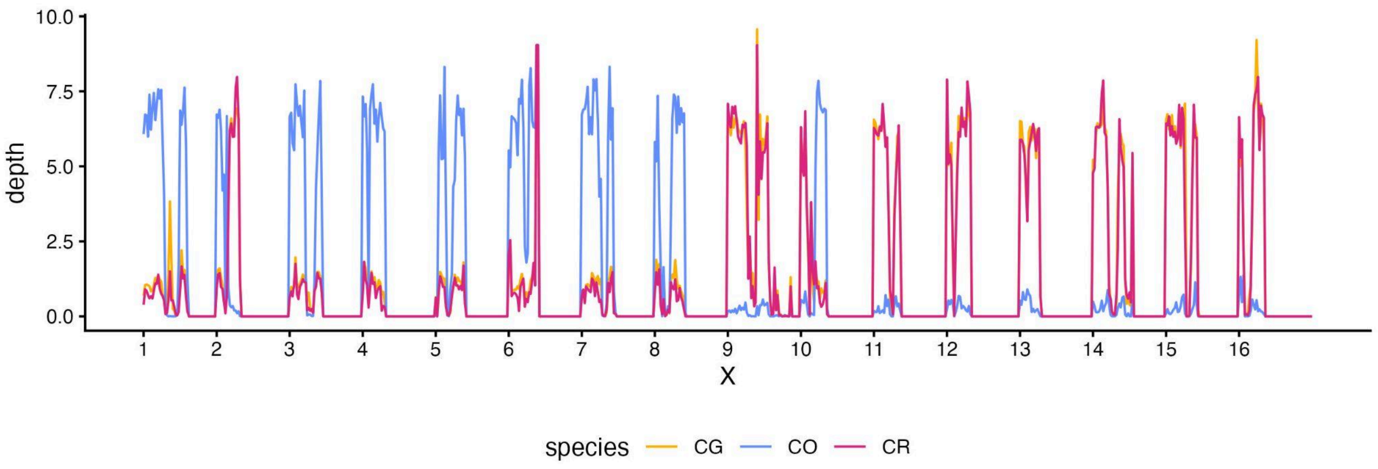


Supplemental Figure 2: Depth analysis of C. rubella, C. grandiflora, and C. orientalis mapped to our reference genome assembly in windows of 1 megabase. Alignment of diploids shows a balanced section of homeologous exchange between Chromosome 10 and Chromosome 2.
Alt Text: colorful line graph showing depth across our reference genome. Depth between C. rubella and C. grandiflora are frequently co-linear and have more coverage on Chromosomes 9 through 16. C. orientalis has higher coverage on Chromosomes 1 through 8. At chromosome 10 around 10000000 basepair, coverage from C. orientalis increases while coverage for C. rubella and C. grandiflora decreases. At Chromosome 2, coverage for C. orientalis decreases while coverage for C. rubella and C. grandiflora increases.

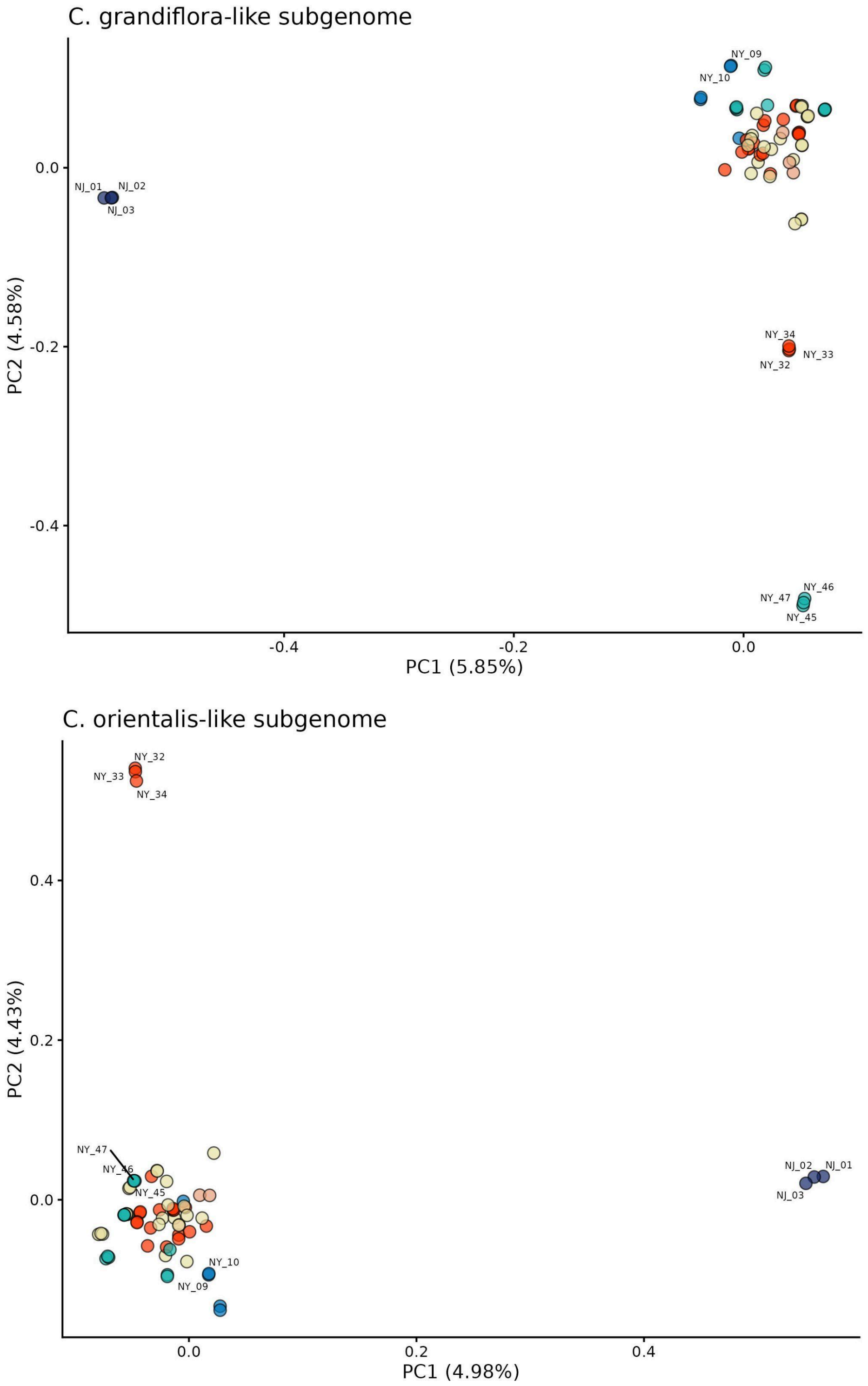


Supplemental Figure 3: genetic principal components 1 and 2 of NYC *C. bursa-pastoris* on the C. grandiflora-like subgenome (top) and the C. orientalis-like subgenome (bottom)
Alt text: colorful scatterplot showing genomic principal components 1 versus 2.

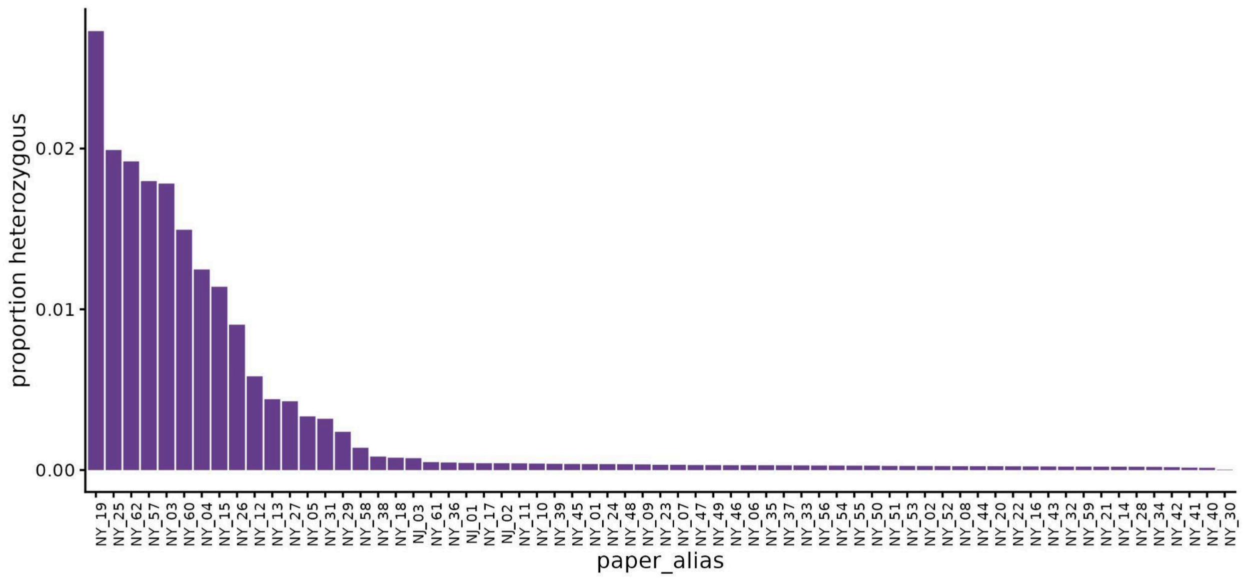


Supplemental figure 4: Individual heterozygosity for NYC metropolitan samples
Alt text: Purple bar graph indicating proportion of heterozygous calls for individuals in the New York City metropolitan area *Capsella bursa-pastoris*. Individuals on the horizontal are ordered by decreasing heterozygosity. A majority of individuals have a heterozygous call proportion below 0.004. Individual NY_19 has notably the highest heterozygous call proportion of 0.0273.

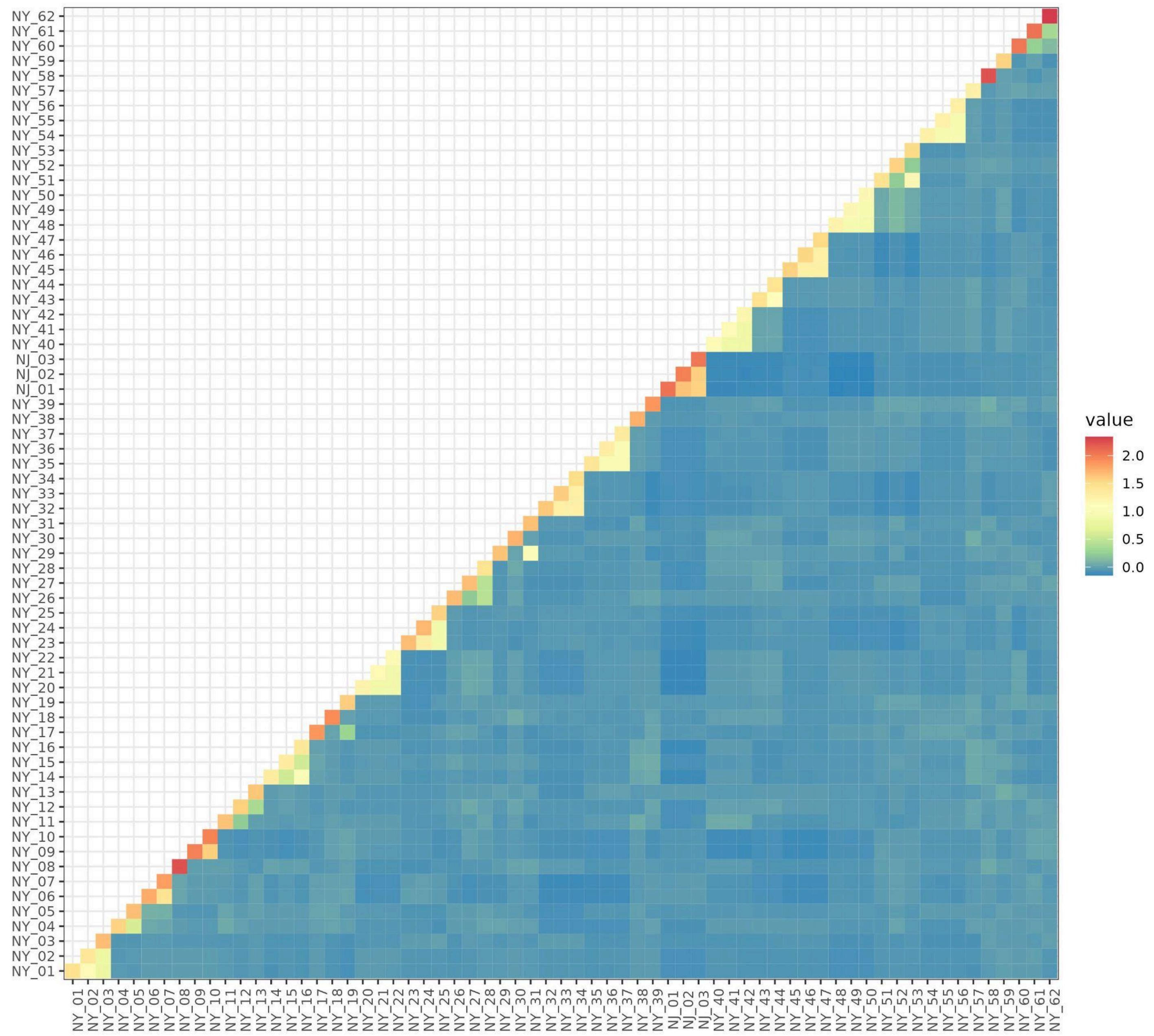


Supplemental figure 5: Relatedness matrix of all New York City metropolitan genotypes.
Alt text: Colorful heatmap showing pairwise relationships between individual samples of New York CIty metropolitan area *Capsella bursa-pastoris*.

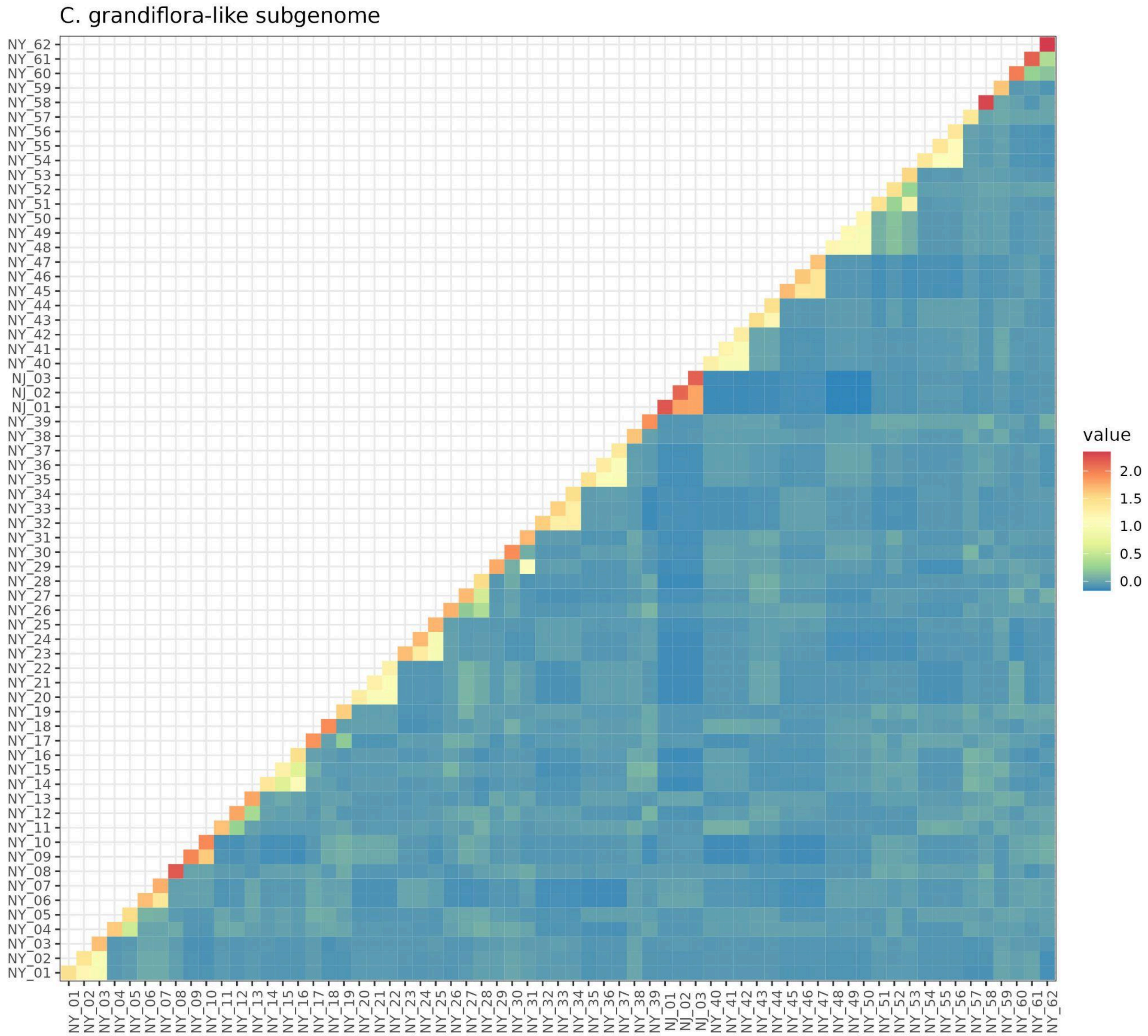


Supplemental figure 6: Relatedness matrix of all New York City metropolitan genotypes on the C. grandiflora-like subgenome.

Alt text: Colorful heatmap showing pairwise relationships between individual samples of New York CIty metropolitan area *Capsella bursa-pastoris*.

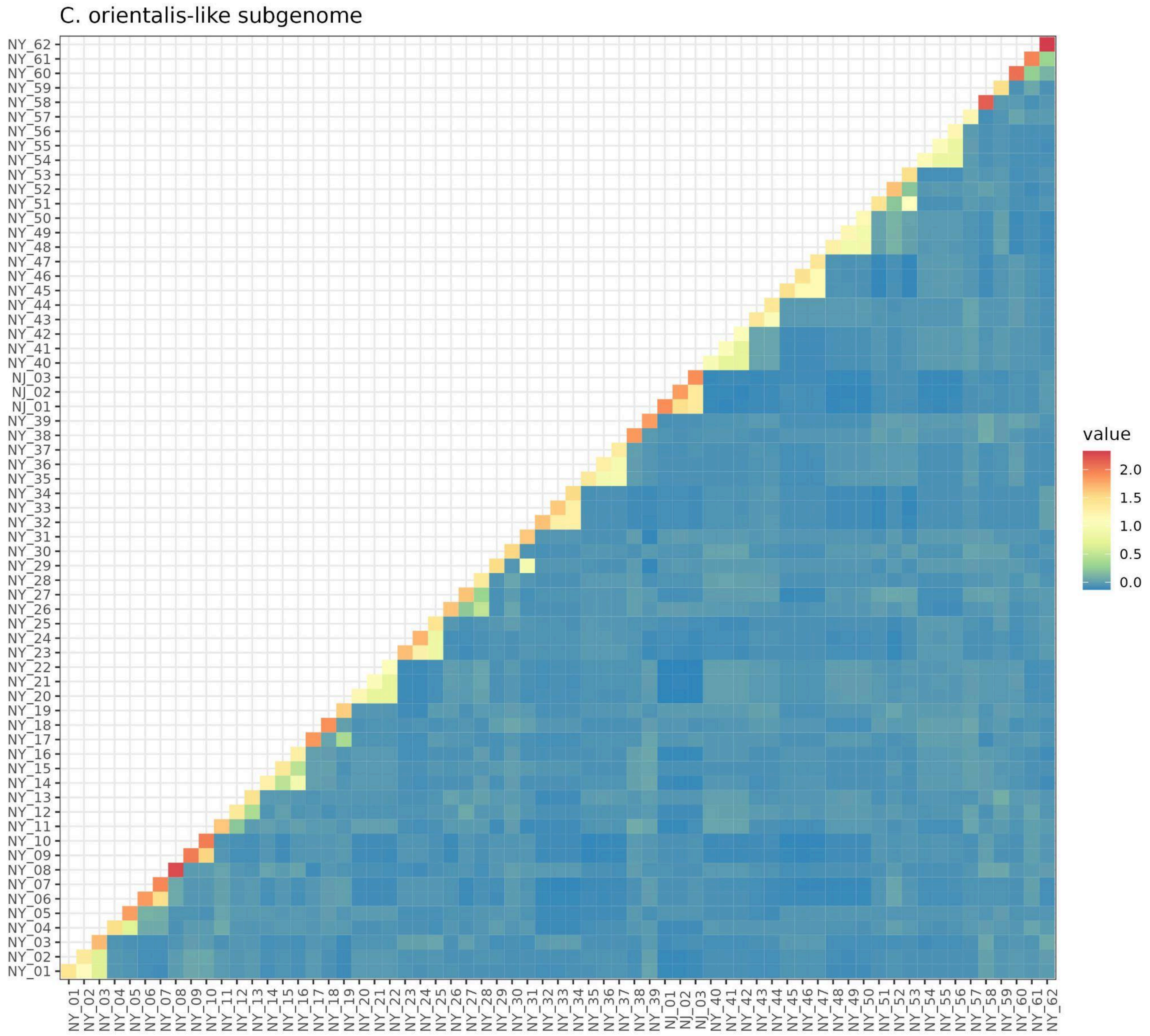


Supplemental figure 7: Relatedness matrix of all New York City metropolitan genotypes on the C. orientalis-like subgenome.

Alt text: Colorful heatmap showing pairwise relationships between individual samples of New York CIty metropolitan area *Capsella bursa-pastoris*.

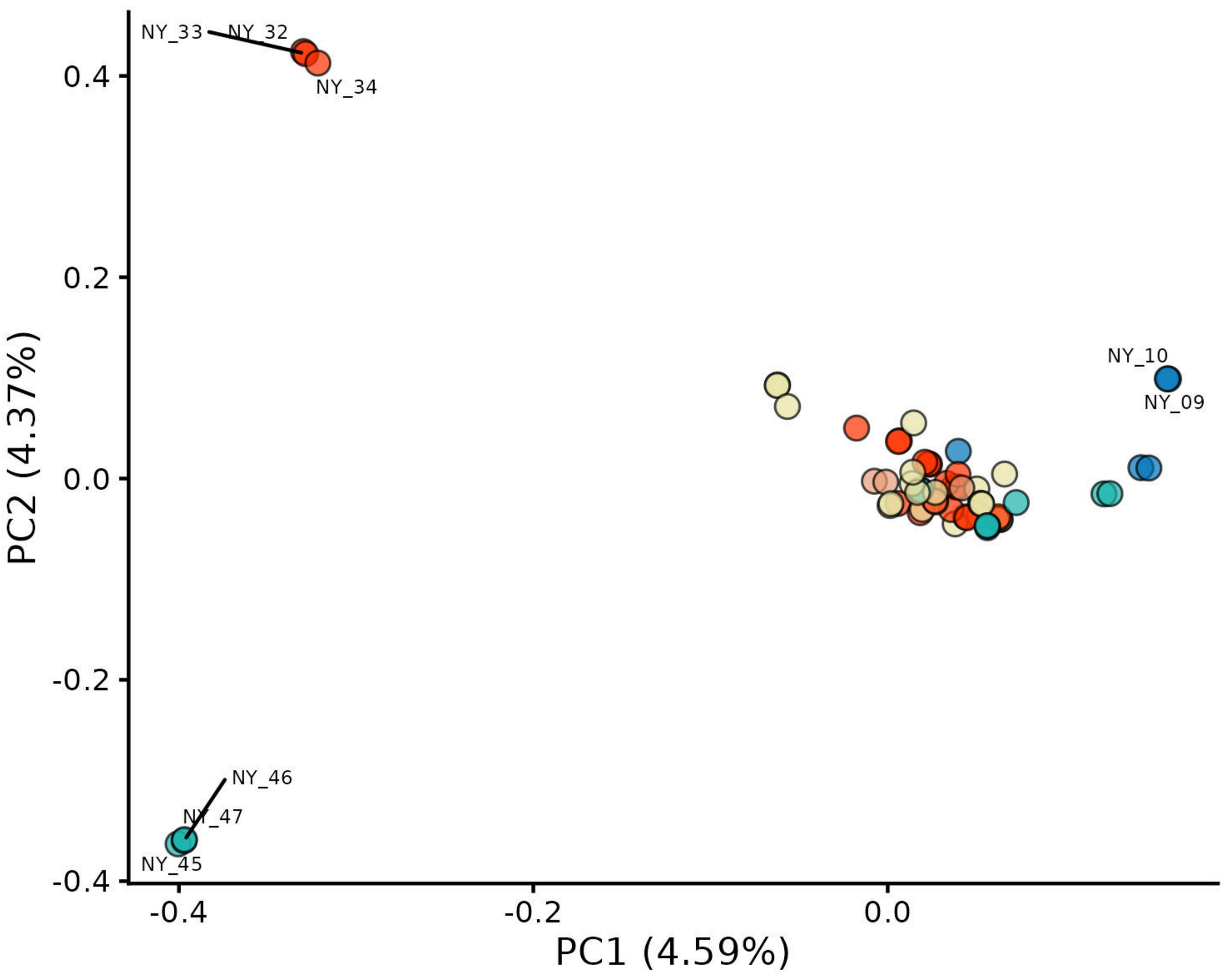


Supplemental Figure 8: genetic principal components 1 and 2 of *C. bursa-pastoris* only within New York State.
Alt text: colorful scatterplot showing genomic principal components 1 versus 2

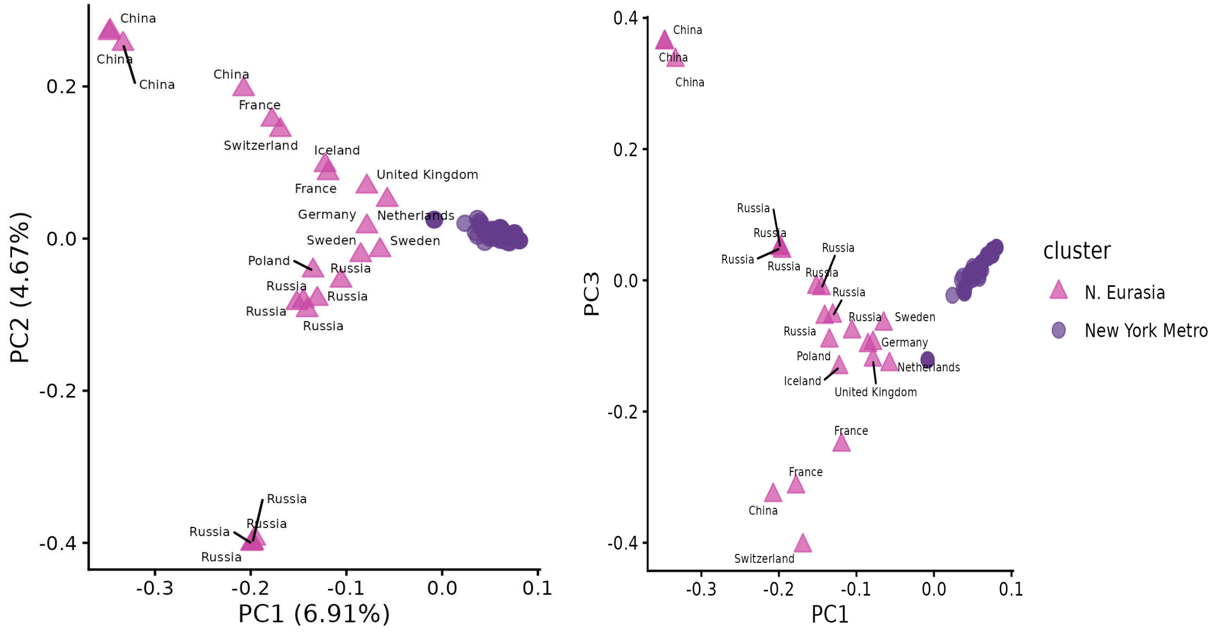


Supplemental Figure 9: Principal component analysis of N. Eurasian ancestry group with NYC metropolitan area. *C. bursa-pastoris* outside of the United States (triangles) are labelled with the country of collection. Principal components 1 and 2 are shown on the left; Principal components 1 and 3 are show on the right panel of the figure.

Alt text: Pink and purple scatterplot showing genomic principal components

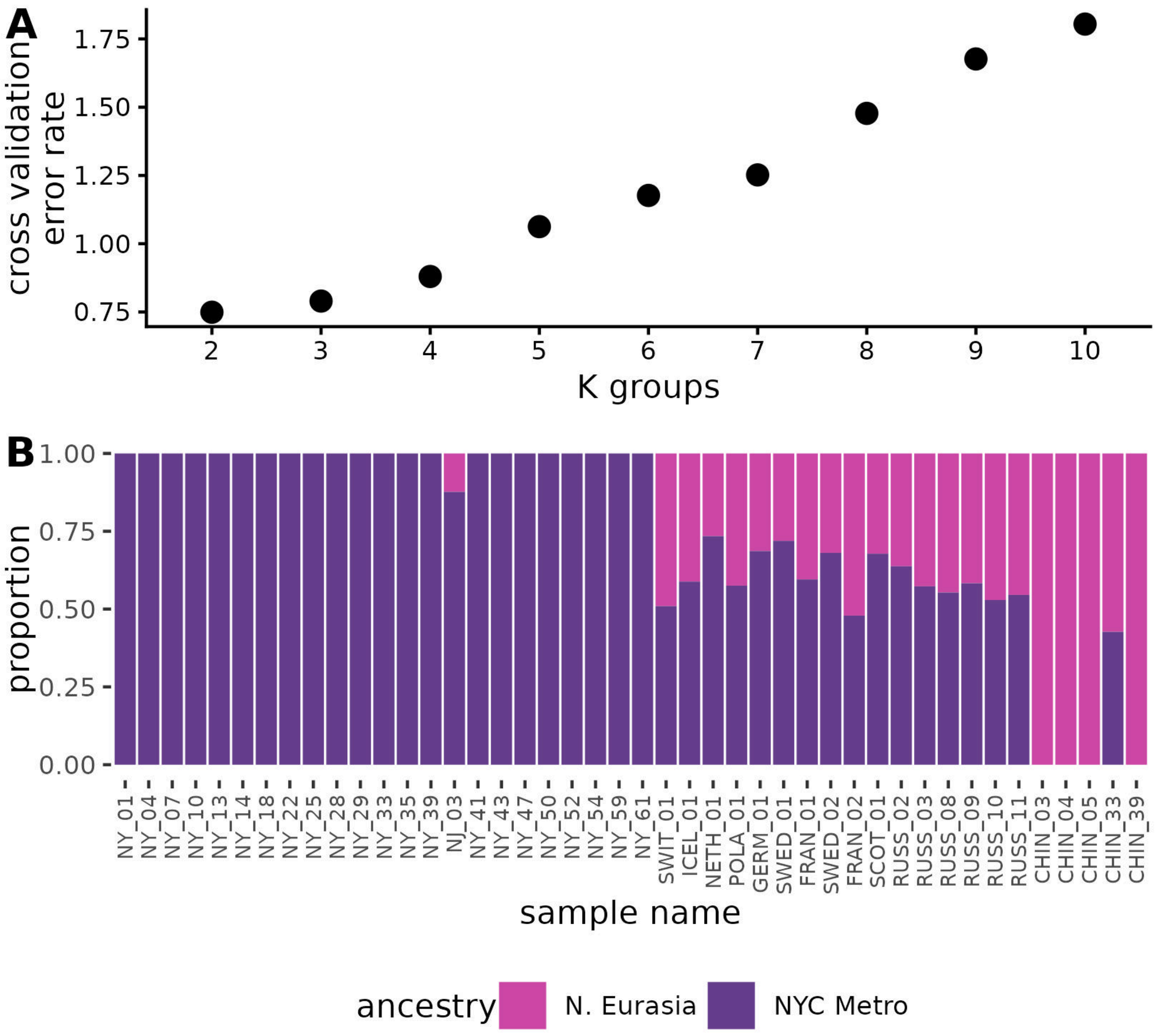


Supplemental figure 10: K-means clustering among N. Eurasian ancestral group and NYC metro *C. bursa-pastoris*. NYC samples have been thinned to 1 per sampling location, and identical Russian CBP have been omitted A) Cross validation error rates of k-groups 2 through 10. B) Ancestry proportion assignment of subsample of NYC metro *C. bursa-pastoris* and N. Eurasian *C. bursa-pastoris* from outside of the United States of America.

Alt text: ADMIXTURE results showing Northern Eurasian *Capsella bursa-pastoris* in the ancestral range can be distinguished from *Capsella bursa-pastoris* in the New York City metropolitan area and K=2 has the lowest cross-validation error rate. Individuals NJ_03, SWIT_01, ICEL_01, NETH_01, POLA_01, GERM_01, SWED_01, FRAN_01, SCOT_01, RUSS_02, RUSS_03, RUSS_08, RUSS_09, RUSS_10, RUSS_11, and CHIN_33 show intermediate ancestry proportions between ancestral Northern Eurasian and NYC ancestry groups

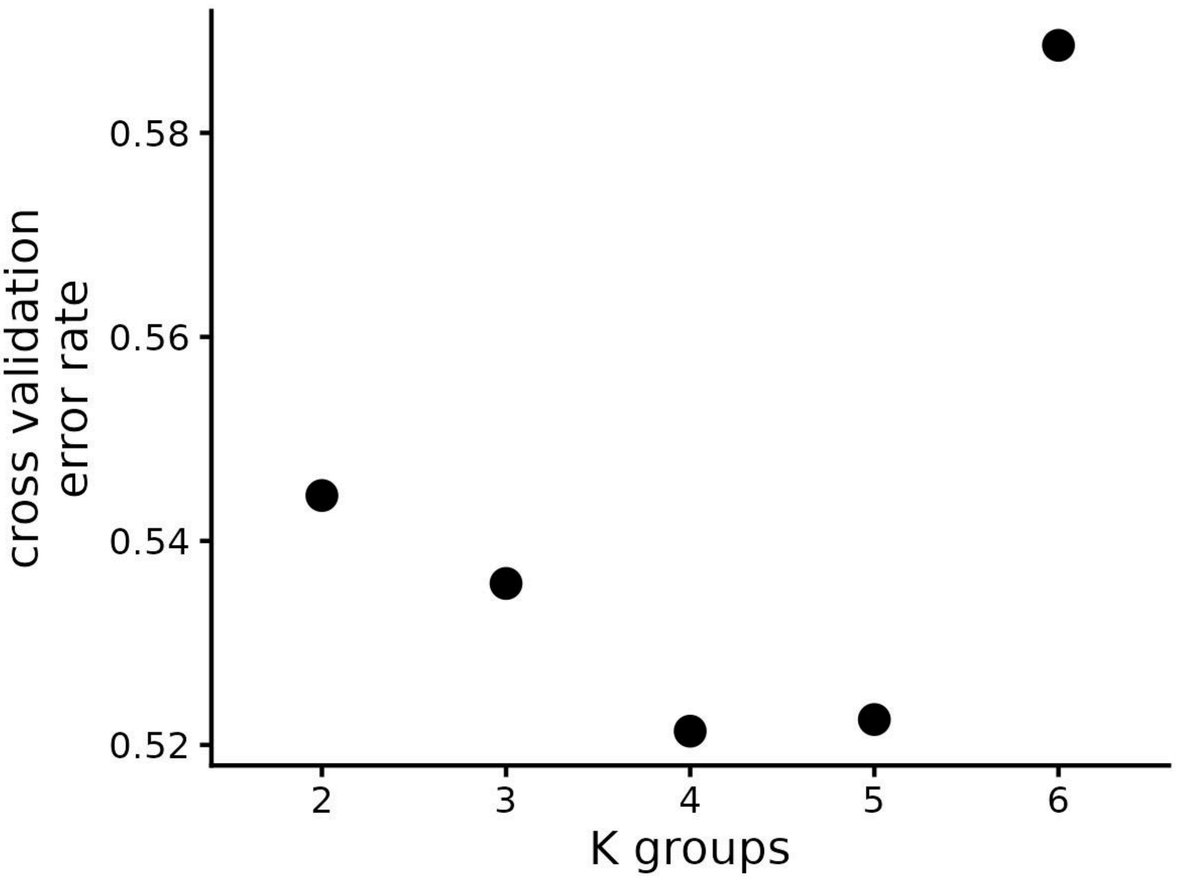


Supplemental figure 11: Cross validation error rates for k-means clustering in ADMIXTURE among ancestral range *C. bursa-pastoris*

Alt text: Black and white scatterplot showing K=4 has the lowest cross-validation error rate

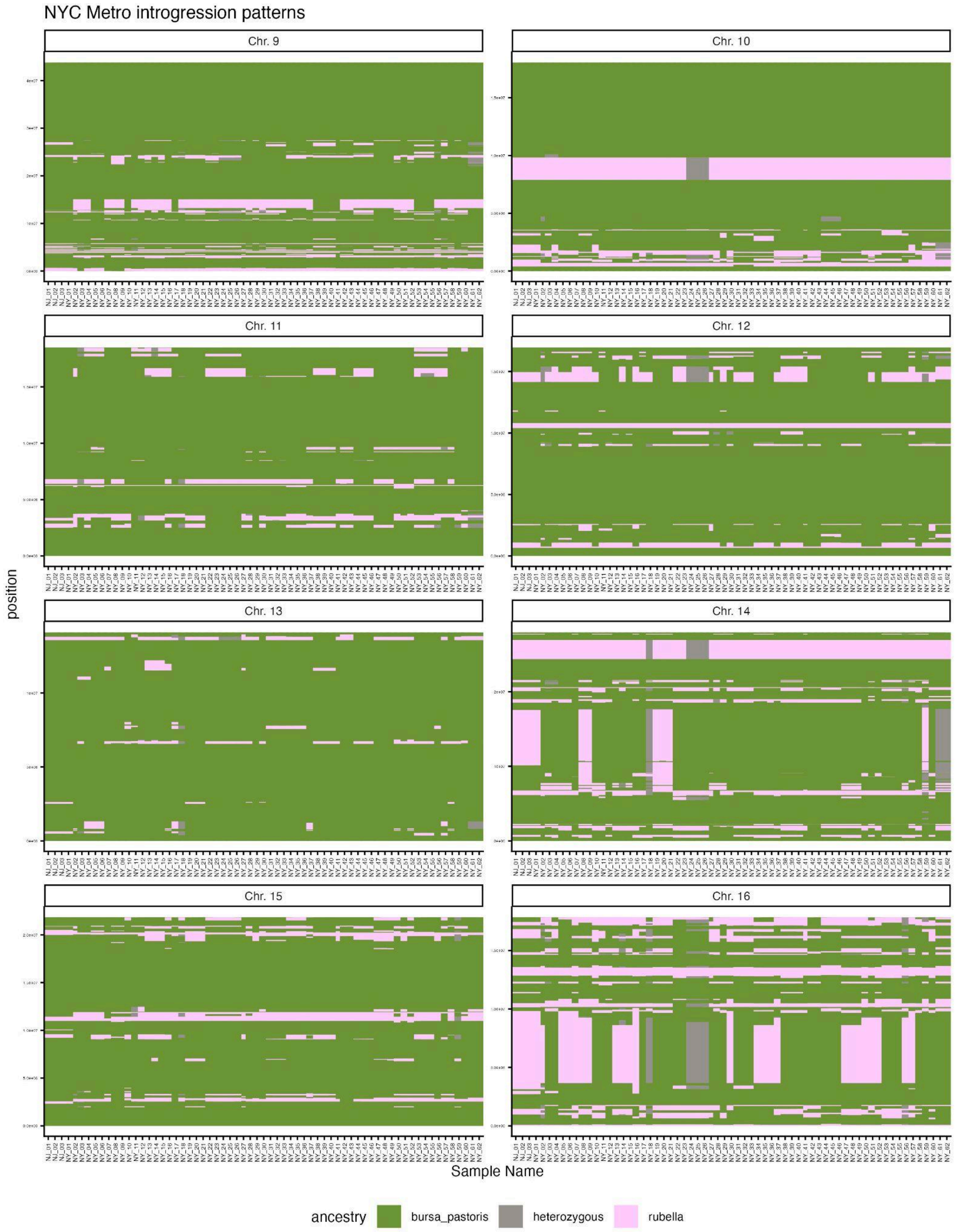


Supplemental figure 12: Inferred ancestry across chromosomes within NYC metro *C. bursa-pastoris*

Alt text: Detailed figure showing inferred ancestry of every NYC *Capsella bursa-pastoris* individual with a panel for each chromosome. *Capsella rubella* ancestry is indicated in pink, *C. bursa-pastoris* ancestry is in green, and heterozygous ancestry in grey.

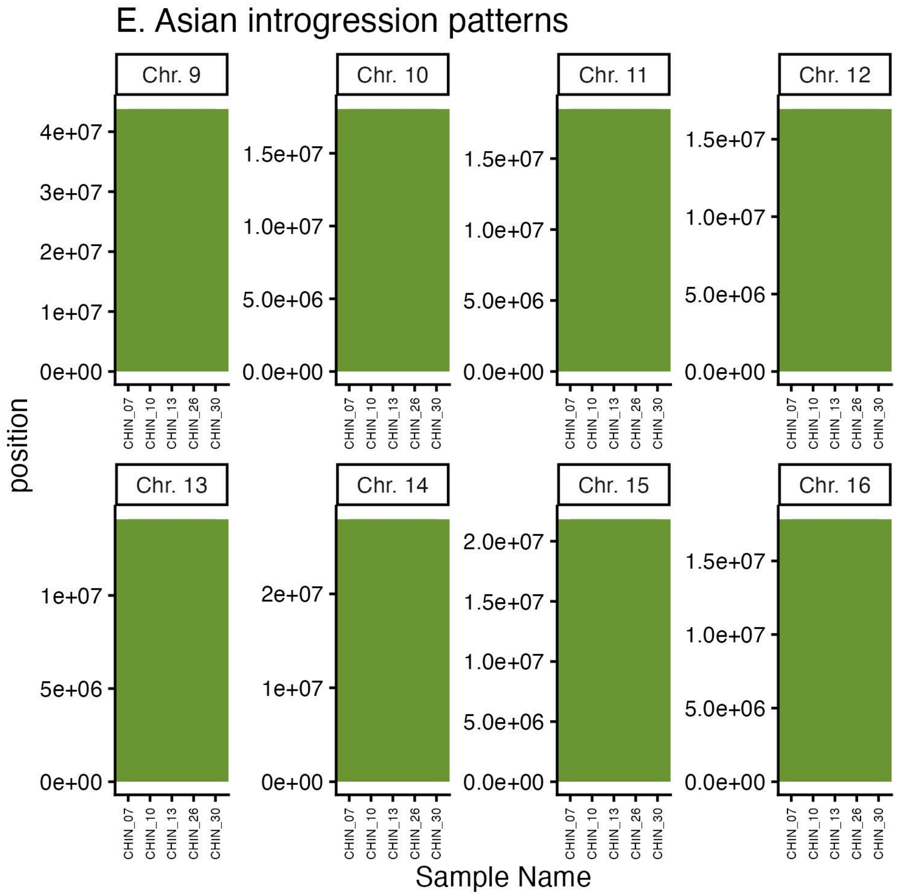


Supplemental figure 13: Inferred ancestry across chromosomes for five randomly selected East Asian C. bursa-pastoris.

Alt text: Detailed figure showing inferred ancestry of subset of East Asian *Capsella bursa-pastoris* individuals with a panel for each chromosome. All individuals show only *C. bursa-pastoris* ancestry across the genome

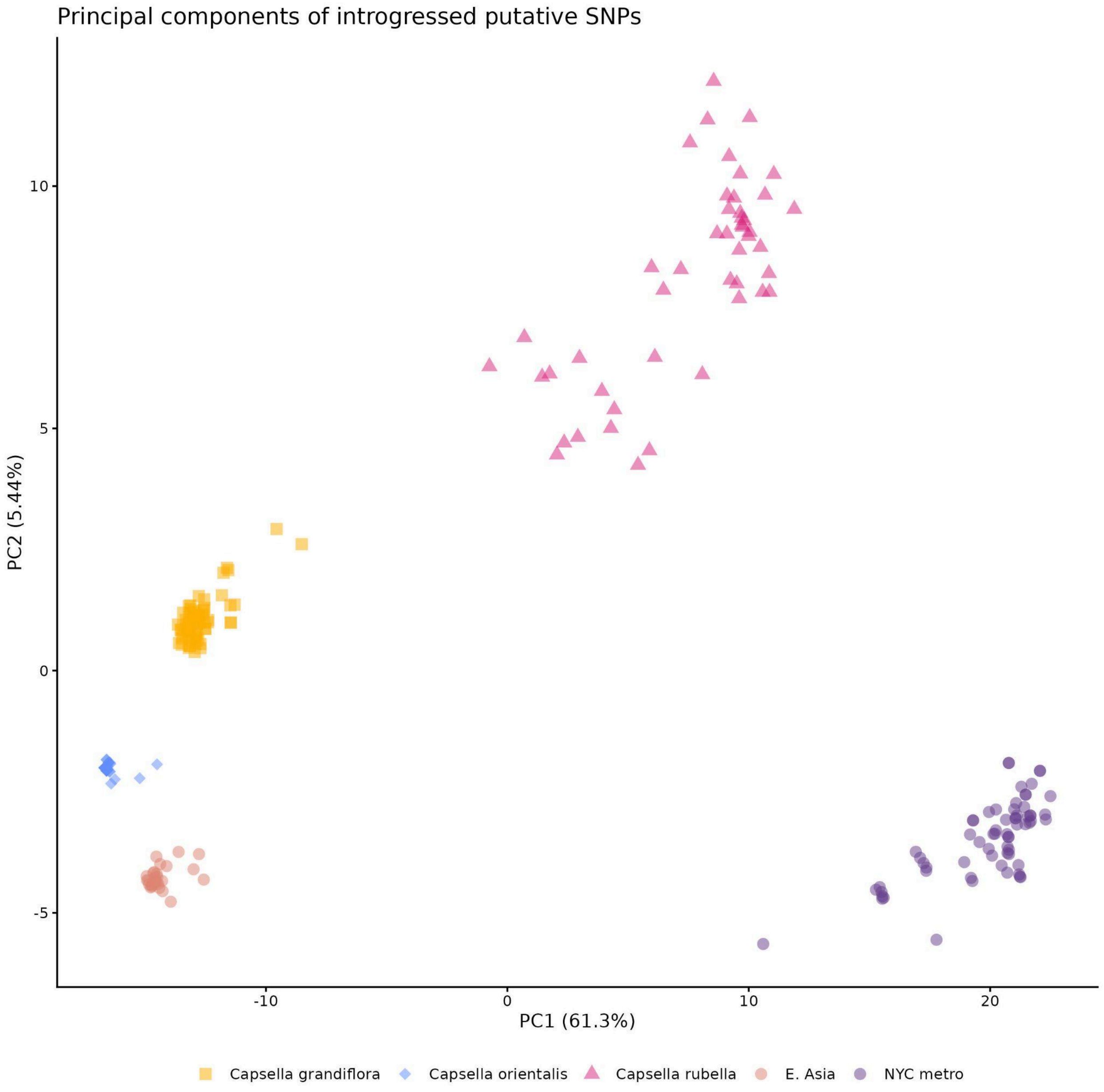


Supplemental figure 14: First and second genetic principal components for NYC metropolitan *Capsella bursa-pastoris* (purple circles), East Asian *C. bursa-pastoris* (orange circles), *C. grandiflora* (yellow squares), *C. rubella* (pink triangles), and *C. orientalis* (blue diamonds) at putatively introgressed *C. rubella* SNPs.

Alt text: Colorful scatter plot showing relationships between *Capsella* species. *Capsella* populations are indicated by shape and color. Principal component 1 on the horizontal axis explains 61.3 percent of the variance in the data and principal component 2 on the vertical axis explains 5.44 percent of the variance in the data. All groups cluster amongst themselves. PC1 separates groups by ancestry where PC2 separates the species from one another.

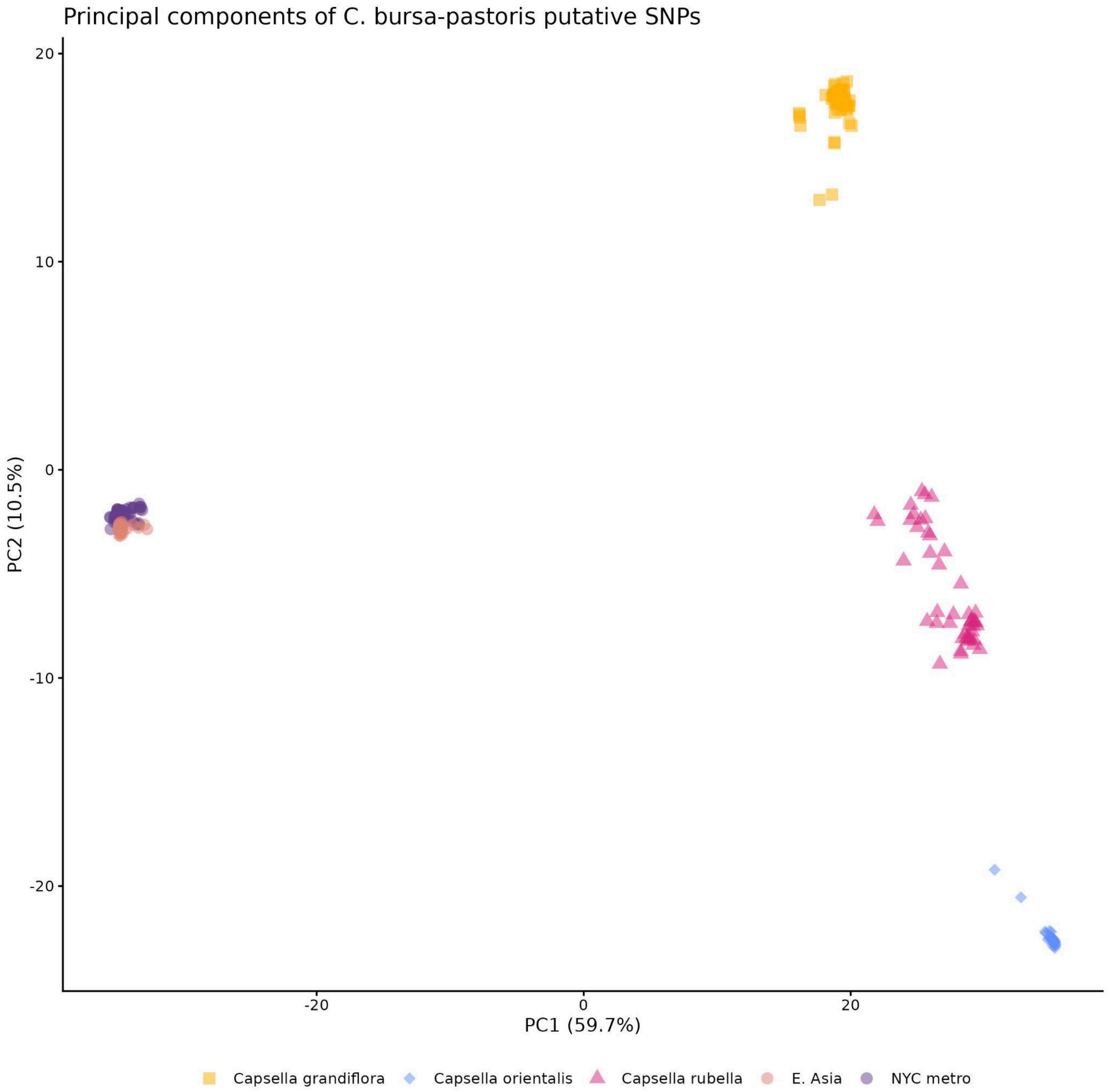


Supplemental figure 15: First and second genetic principal components for NYC metropolitan *Capsella bursa-pastoris* (purple circles), East Asian *C. bursa-pastoris* (orange circles), *C. grandiflora* (yellow squares), *C. rubella* (pink triangles), and *C. orientalis* (blue diamonds) at putatively non-introgressed SNPs.
Alt text: Colorful scatter plot showing relationships between *Capsella* species. *Capsella* populations are indicated by shape and color. Principal component 1 on the horizontal axis explains 59.7 percent of the variance in the data and principal component 2 on the vertical axis explains 10.5 percent of the variance in the data. NYC *C. bursa-pastoris* and East Asian *C. bursa-pastoris* cluster on top of one another.
